# CRISPR-Cas9 for selective targeting of somatic mutations in pancreatic cancers

**DOI:** 10.1101/2023.04.15.537042

**Authors:** Selina Shiqing K. Teh, Kirsten Bowland, Alexis Bennett, Eitan Halper-Stromberg, Alyza Skaist, Jacqueline Tang, Fidel Cai, Antonella Macoretta, Hong Liang, Hirohiko Kamiyama, Sarah Wheelan, Ming-Tseh Lin, Ralph H. Hruban, Robert B. Scharpf, Nicholas J. Roberts, James R. Eshleman

**Author notes:** Corresponding author: James R. Eshleman, MD, PhD. Department of Pathology and Oncology, 1550 Orleans Street, Suite 344, Baltimore, MD, 21231. These authors contributed equally to this work (Selina Shiqing K. Teh, Kirsten Bowland).

## Abstract

Somatic mutations are desirable targets for selective elimination of cancer, yet most are found within the noncoding regions. We propose a novel, cancer-specific killing approach using CRISPR-Cas9 which exploits the requirement of a protospacer adjacent motif (PAM) for Cas9 activity. Through whole genome sequencing (WGS) of paired tumor minus normal (T-N) samples from three pancreatic cancer patients (Panc480, Panc504, and Panc1002), we identified an average of 417 somatic PAMs per tumor produced from single base substitutions. We analyzed 591 paired T-N samples from The International Cancer Genome Consortium and discovered medians of ∼455 somatic PAMs per tumor in pancreatic, ∼2800 in lung, and ∼3200 in esophageal cancer cohorts. Finally, we demonstrated >80% selective cell death of two targeted pancreatic cancer cell lines in co-cultures using 4-9 sgRNAs, targeting noncoding regions, designed from the somatic PAM discovery approach. We also showed no off-target activity from these tumor-specific sgRNAs through WGS.

**Statement of significance:** This study demonstrates the potential of CRISPR-Cas9 as a novel and selective anti-cancer strategy. It requires just a few targets to induce double strand breaks for significant cytotoxicity. Our findings markedly expand the repertoire of targetable mutations in cancers and support genetically targeting other adult solid tumor types.

## Introduction

Somatic mutations accumulate as we age and are clonally propagated from the cancer initiating cell to all neoplastic daughter cells (1). These somatic mutations genetically define the malignant cell population and are exploited as therapeutic targets. Indeed, most targeted therapies focus on mutations within coding regions, as drugs and vaccines have been designed to target the mutated proteins or produce synthetic lethality. The targeted mutations are commonly driver mutations or mutations with known roles in carcinogenesis, inevitably limiting the number of targetable mutations available (2). Meanwhile, most mutations in cancers are found within noncoding regions which make up 98% of the human genome (3). Most of these tumor-specific “passenger mutations” are traditionally thought to be inconsequential to cell fitness or haven’t had their functions elucidated, and therefore lacking therapeutic value (4). This prompted us to develop a strategy to turn this vast number of mutations into targets of therapeutic importance, independent of their individual genetic functions.

CRISPR-Cas9 is a programmable endonuclease or “molecular scissor” (5–7). It can be designed to induce double strand breaks (DSBs) at desired locations in the human genome, and in theory, can be highly selective. Several papers have reported cytotoxicity associated with multiple Cas9-induced DSBs (8,9), and have either developed various methods to circumvent the toxicity to increase the success rate of gene editing (10–12), or exploited this toxicity to achieve cell-specific killing (13,14). However, CRISPR-Cas9 is known to tolerate mismatches at the single-guide RNA (sgRNA) target regions, contributing to off-target effects (15–18).

Our approach is inspired by the bacterial adaptive immune system, in which protospacer adjacent motifs (PAMs) have been evolutionarily selected to differentiate between host and viral DNA, that otherwise contain the exact same sequence, to selectively eliminate the pathogen while leaving the bacterial host intact (19). The 3-nucleotide 5’-NGG-3’ PAM sequence, recognized by *S. pyogenes* Cas9, serves as a binding signal, for which Cas9 neither binds nor cuts the target in its absence (20,21), significantly decreasing the risk of off-targeting. We therefore determined the type of somatic mutations in cancers that would create the largest number of novel CRISPR-Cas9 target sites and developed a straightforward bioinformatics pipeline to identify single base substitutions (SBSs) that formed novel, cancer-specific PAMs. We identified hundreds to thousands of somatic PAMs in different tumor types, and these somatic PAMs could serve as cancer-specific targets to selectively kill the malignant cell population.

## Results

### Development of PAM discovery approach

We tested two approaches with the potential leading to highly selective target cell killing with minimal off-target risk. As pancreatic cancer (PC) is one of the most lethal cancers with a dismal five-year survival rate of only 11.5% (22), we chose PC as the model to test our hypothesis. We previously generated primary PC cell lines and their corresponding normal cell lines from three PC patients (Panc480, Panc504, and Panc1002; table S1) (23) and performed whole genome sequencing (WGS). We then performed tumor-normal (T-N) subtraction to identify somatic mutations. All three PC samples harbored deleterious mutations in *KRAS*, *CDKN2A*, *SMAD4*, and *TP53*, which are the most common driver mutations in PCs (table S1).

We first considered structural variants (SVs) as they could juxtapose a new target DNA sequence next to an existing NGG PAM (figure S1A-B). This could theoretically decrease the risk of off-target effects, as the resulting breakpoint is significantly different from the original sequence (figure S1C). We discovered an average of 35 SVs per cell line by comparing tumor to normal, and validated 84.9% of them by PCR amplification across the breakpoint and Sanger sequencing (table S2, figure S1C). We found an average of 22 novel SVs juxtaposed next to an existing PAM per cell line (table S2). Using our sgRNA selection criteria (see Supplementary methods), we obtained an average of 17 sgRNAs per cell line with minimal off-targeting risk (table S2).

Next, we attempted to discover novel PAMs created from SBSs (Figure 1A-B). A somatic NGG PAM can arise through a SBS that creates a novel G from A/T/C, and this novel G is adjacent to an existing G immediately upstream or downstream of the new G (Figure 1A-B). The same concept applies to the complementary strand (CCN). Our PC samples harbored mutational signatures that produced novel Cs and Gs (Figure 1C), with the most common signatures being SBS1, 5, and 40 (figure S2). These were all clock-like signatures (24–26), suggesting that aging itself gave rise to novel PAMs. We then developed a program, PAMfinder, to discover somatic base substitutions that produced novel PAMs in a given tumor sample.

**Fig. 1.**
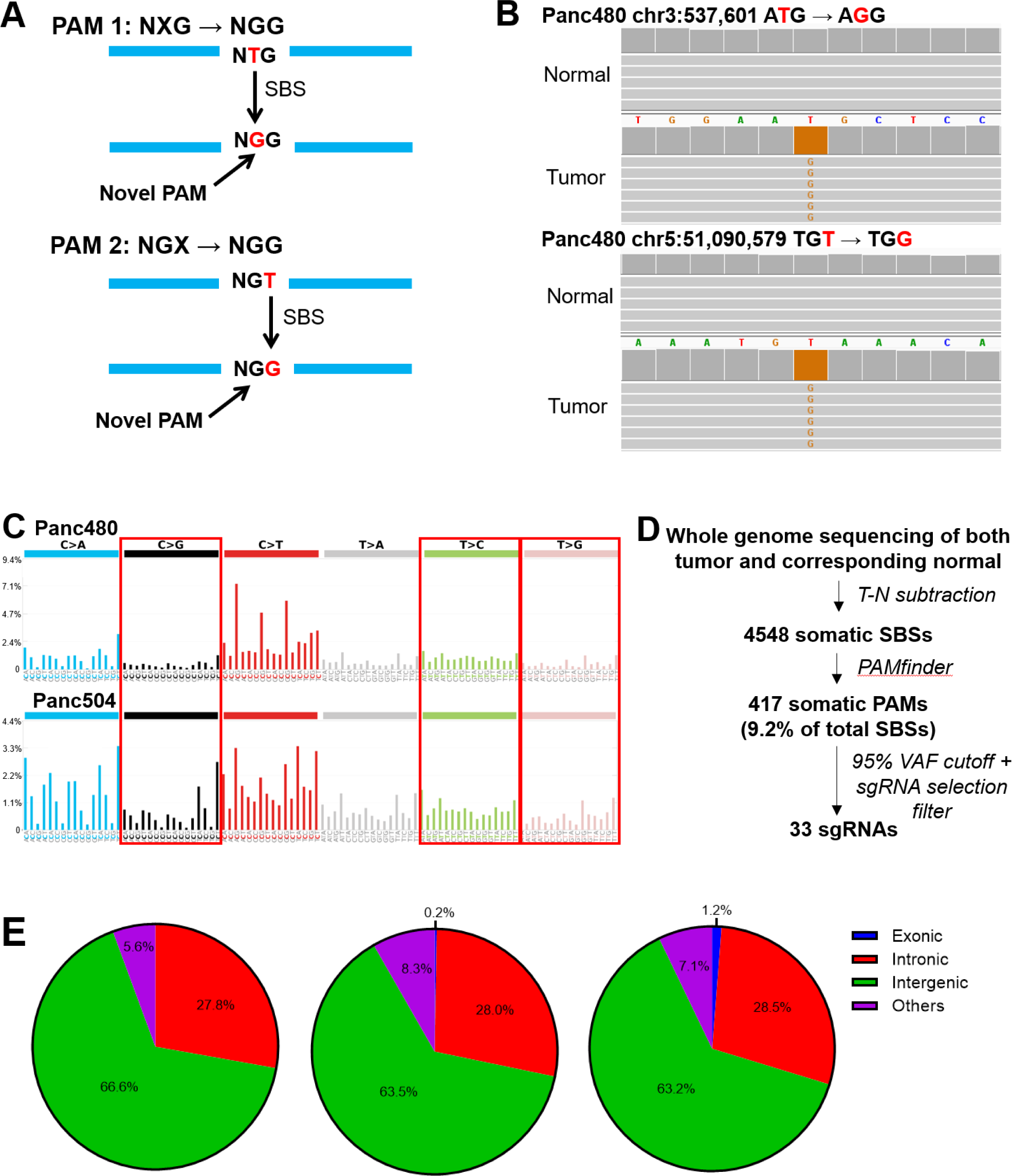
Somatic PAM discovery yielded hundreds of novel PAMs in pancreatic cancers (PCs). (A) A somatic NGG protospacer adjacent motif (PAM) can arise through a single base substitution (SBS) that creates a novel G from A/T/C (indicated as X), and this novel G is adjacent to an existing G immediately downstream (PAM 1) or upstream (PAM 2) of the novel G. Examples of T>G are shown. (B) Two somatic PAMs in the Panc480 tumor were both absent in their corresponding normal tissues. (C) Mutational signatures of two PCs, Panc480 and Panc504. The proportions of SBS creating novel Gs and Cs that could potentially form novel PAMs were highlighted in red boxes. Y-axis is the percentage of SBS. (D) Workflow of somatic PAM discovery. (E) Proportions of novel PAMs discovered in Panc480 (left), Panc504 (middle), and Panc1002 (right) that were located in different regions of the human genome. Others included non-coding RNAs, untranslated regions, and 1kb regions upstream/downstream of transcription start/end sites. For Panc480, no novel PAMs were found in exons.

We identified an average of 4,548 SBSs per sample, of which 9.2% created somatic PAMs (mean=417; Figure 1D, table S3). A variant allele frequency (VAF) cutoff of 30% was used to exclude mutations that might be subclonal or have arisen through *in vitro* culture of these cell lines. We selected novel PAMs with VAFs >95% (mean=63) for initial functional testing of sgRNAs as targeting them should produce the highest toxicity. Of these, we were able to design an average of 33 sgRNAs, one for each mutation, that had minimal risk for off-target activity (see Methods, Figure 1D, table S3). We confirmed all qualifying mutations, except two, through Sanger sequencing (table S3). A similar approach using whole exome sequencing (WES) data failed to yield sufficient targets (mean=1; table S4). This was because the majority of cancer-specific PAMs were located in noncoding regions, with 64.4% of somatic PAMs located in intergenic regions, 28.1% in introns, 0.5% in exons, and the remaining 7.0% in other regions such as noncoding RNAs (Figure 1E). Thus, we concluded that the WGS-based PAM discovery approach using SBSs was more productive than the SV and WES approaches, and provided hundreds of novel PAMs per cancer as potential CRISPR-Cas9 target sites.

### High prevalence of novel PAMs in different tumor types

To determine the prevalence of somatic PAMs in different tumor types, we obtained T-N variant call files (VCFs) from the ICGC Data Portal, including pancreatic (APGI-AU and PACA-CA), lung (LUCA-KR), and esophageal cancer (OCCAMS-GB) samples (27). We performed tumor purity correction and analyzed the data using PAMfinder (Figure 2A, table S5). Overall, we found that the number of base substitutions and number of somatic PAMs from the two PC cohorts, APGI-AU (N=44) and PACA-CA (N=130), were comparable to findings from our discovery PC lines, in which a median of 478.5 and 430.5 somatic PAMs per tumor were identified, respectively (Figure 2B-C, table S3 & S5). Regarding the 29 lung cancer samples and 388 esophageal cancer samples, the number of PAMs identified was >5 fold higher than that of PCs, with medians of 2790 and 3235.5 somatic PAMs per tumor, respectively (Figure 2C, table S5). Since the number of base substitutions were also higher in lung cancers (median=30553) and esophageal cancers (median=20106) compared to PCs (median=5890.5 and 5354.5), these results suggested that tissue-specific factors, such as environmental mutagens, contributed to the varying number of mutations present (Figure 2B, table S5).

**Fig. 2.**
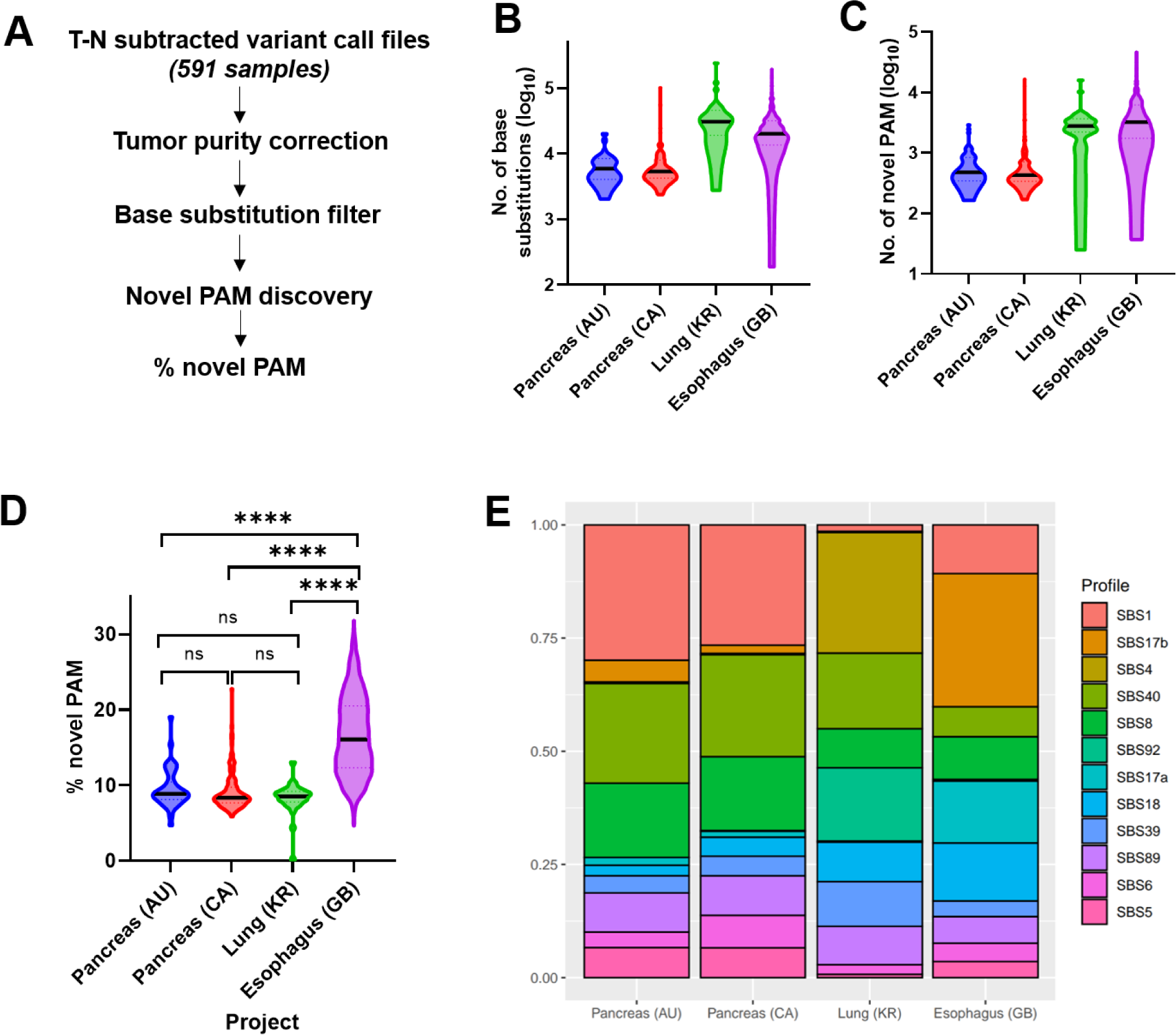
Hundreds to thousands of somatic PAMs were found in different adult solid tumor types. (A) Workflow of PAM discovery in 591 tumor samples using T-N subtracted variant call files from the ICGC Data Portal (60). All analyses were corrected based on the tumor purity of individual sample. Samples from four cohorts were included: APGI-AU (Pancreas (AU); N=44), PACA-CA (Pancreas (CA); N=130), LUCA-KR (Lung (KR); N=29), and OCCAMS-GB (Esophagus (GB); N=388). (B-C) Truncated violin plots present the total number of (B) base substitutions and (C) novel PAMs in each cohort. (D) Truncated violin plot presents the percentage of base substitutions that created somatic PAMs. Kolmogorov-Smirnov tests; ns indicates non-significant, **** indicates *P<0.0001*. (E) Mutational spectra analysis in each cohort.

While the proportions of base substitutions that gave rise to somatic PAMs (% novel PAM) were similar among PCs (median=8.9% and 8.4%) and lung cancers (median=8.5%), esophageal cancers had significantly higher % novel PAM at 16.1% (interquartile range=12.3-20.5%; *P<0.0001*; Figure 2D, table S5). To investigate the potential mechanism contributing to the higher % novel PAM, we performed mutational signature analysis on all samples. We found that the two sets of PC samples showed similar top ranked mutational signatures that were consistent with our discovery PC cell lines (SBS1 and SBS40; Figure 2E, figure S2). The top mutational signatures for lung cancers, SBS4 and SBS92, were associated with tobacco smoking, in which SBS92 had a predominant T>C mutation signature (Figure 2E) (25,28,29). Notably, the top ranked mutational signature of esophageal cancers, SBS17b, distinguished itself from the other tumor types (Figure 2E), consistent with studies published with these samples (30,31). It was characterized primarily by a T>G transversion with an unknown etiology, but previous studies had associated it with fluorouracil (5FU) treatment and possibly damage by reactive oxygen species (30,32). Based on our analyses of different large tumor cohorts, we concluded that somatic base substitutions yielded hundreds, if not thousands, of novel PAMs in each tumor, and these findings were tissue, and potentially, treatment-dependent.

### Selective cell killing with CRISPR-Cas9

To estimate the number of sgRNAs required to generate significant cytotoxicity in PC cells, we designed sgRNAs with increasing number of target sites in the human genome (designated as multitarget sgRNAs), transduced them into two PC cell lines (Panc10.05 and TS0111), and performed clonogenicity assays (table S6). All perfect target sites and potential off-target sites of the multitarget sgRNAs were located in the noncoding regions of the human genome to prevent gene essentiality-linked cytotoxicity from confounding our interpretations. We found that growth inhibition increased with the number of sgRNA target sites (Figure 3A). The 3-target sgRNA, 52F(3), exhibited >75% growth inhibition in both cell lines, suggesting that a few DSBs could produce significant cytotoxicity. The 12- and 14-target sgRNAs, 230F(12) and 164R(14), displayed >95% growth inhibition, comparable to the positive control sgRNAs targeting LINE-1 and Alu elements in the human genome.

**Fig. 3.**
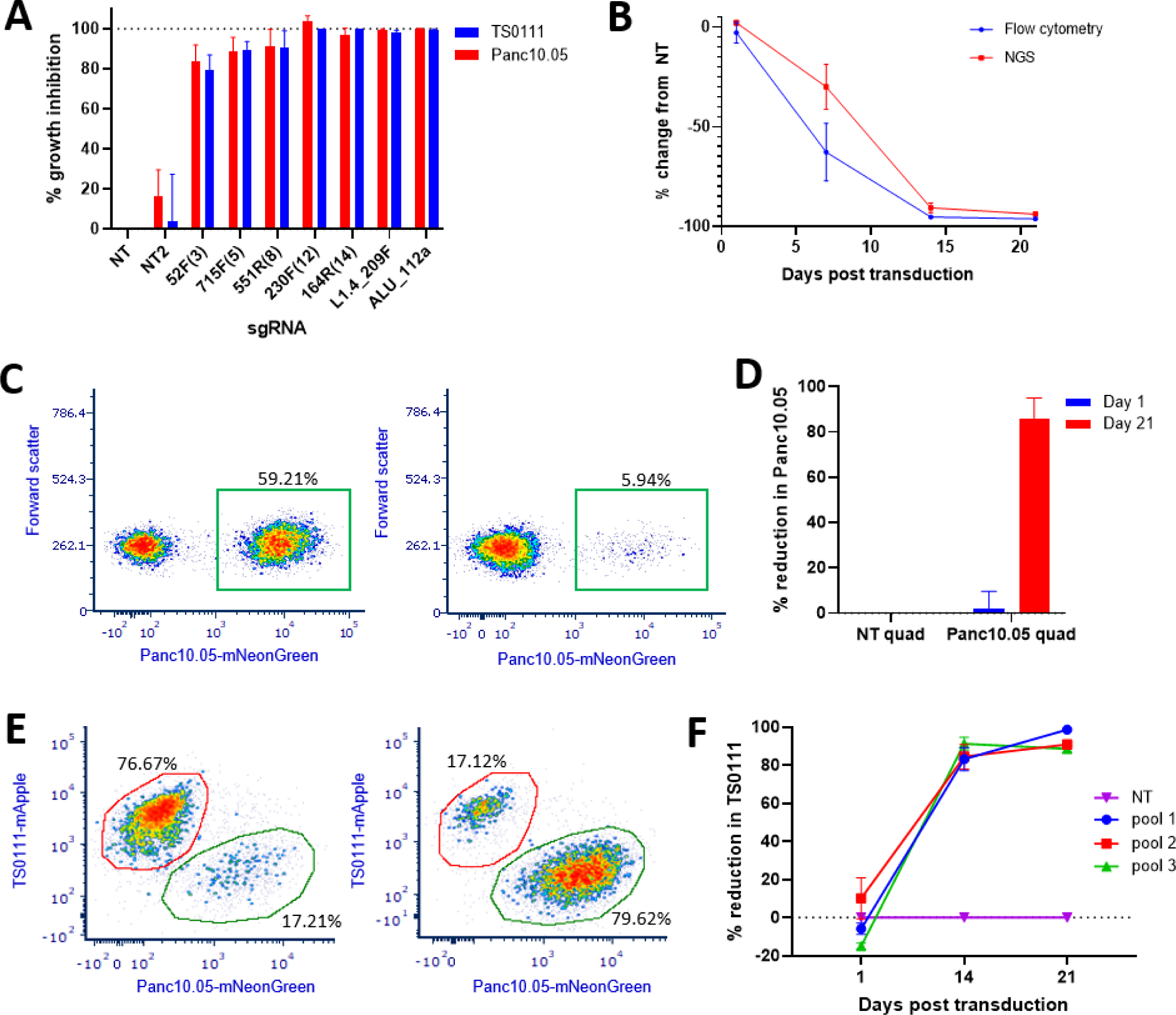
Selective cell killing with low number of sgRNAs designed from our somatic PAM discovery approach. (A) Growth inhibition of two PC cell lines, Panc10.05 and TS0111, treated with non-targeting sgRNAs (NT and NT2), 3-, 5-, 8-, 12-, and 14-target sgRNAs, and repetitive region-targeting sgRNAs (L1.4_209F, ALU_112a). N=3; mean ± SEM are shown. (B) Panc10.05 cell population in co-cultures of Cas9-expressing Panc10.05 and mouse fibroblast (NIH 3T3) cell line transduced with human-specific 230F(12) sgRNA was quantified over time using flow cytometry and a mouse-human NGS assay. N=3; mean ± SEM. (C-D) Co-cultures of Panc10.05 (labeled with mNeonGreen) and TS0111 cell mixtures were transduced with 4-sgRNA expression vectors that included either all non-targeting sgRNAs (NT quad) or Panc10.05-specific sgRNAs (Panc10.05 quad), and flow cytometry was performed to quantify mNeonGreen-positive cells. (C) Flow cytometry analyses of one replicate on day 21 post transduction were shown. Left panel is cells treated with NT quad and the right panel is with Panc10.05 quad. (D) Percentage reductions of Panc10.05 relative to NT quad on day 1 and day 21 post transduction of sgRNAs were shown. N=3; mean ± SEM. (E-F) Co-cultures of TS0111 (labeled with mApple) and Panc10.05 (labeled with mNeonGreen) were treated with three different pools of TS0111-specific sgRNAs (9 sgRNAs per pool) or the equivalent doses of non-targeting sgRNA controls (NT), and flow cytometry was performed to quantify cells that were positive for either mNeonGreen or mApple. (E) Flow cytometry analyses of one replicate on day 14 post transductions were shown. Left panel is cells treated with NT and the right panel is with Pool 1. (F) Percentage reductions of TS0111 relative to NT on day 1, 14, and 21 post transduction were shown. N=3; mean ± SEM.

To demonstrate proof-of-concept selectivity of CRISPR-Cas9, we generated Cas9-expressing mouse (NIH3T3 and Panc02) and human PC (TS0111 and Panc10.05) cell lines with confirmed Cas9 activity (figure S3A-B), established mouse-human cell line co-cultures, and transduced them with a multitarget sgRNA that targets 12 sites in the human genome (230F(12)) but none in the mouse genome (table S7). Using both flow cytometry and a mouse-human next generation sequencing (NGS) assay (figure S3C-D), we saw >95% reduction of the human PC cells in different co-cultures (Figure 3B, figure S3E-F). Human-specific cell killing was dependent on both functional Cas9 and the human-specific sgRNA (figure S3E-G), showing that CRISPR-Cas9 is capable of selectively eliminating cancer cells.

Finally, we tested the hypothesis that we could selectively target a patient’s cancer by treating with sgRNAs designed from our somatic PAM discovery approach. Using PAMfinder, we identified a list of Panc10.05-specific sgRNAs that had >95% VAF in the cell line and analyzed them to identify sgRNAs that had high cutting efficiency and minimal potential off-target activity (see Methods). We then cloned four of them into a multiplex sgRNA expression vector that expressed four sgRNAs simultaneously (Panc10.05 quad) and, in parallel, cloned four non-targeting sgRNAs into a second multiplex expression vector (NT quad) as negative control (fig. S4A, table S8). The target sites of these Panc10.05-specific sgRNAs were located in either introns or intergenic regions, and deep sequencing showed that these sgRNAs induced mutations in Panc10.05 and not in the non-target cell line (fig. S4B, table S8). We then transduced the quads into co-cultures of Panc10.05 and TS0111, and observed an average of 86% selective reduction of Panc10.05 cells relative to negative control 21 days after transduction (Figure 3C-D). To examine Cas9 selectivity in a second cell line, we chose 27 sgRNA sequences that had >95% VAF in the cell line and cloned each of them into a sgRNA expression vector (table S9). We then transduced three separate pools of sgRNA, 9 sgRNAs per pool (table S9), into Panc10.05-TS0111 co-cultures. We found that >83% selective reduction of TS0111 was achieved by day 14, and >88% by day 21 (Figure 3E-F, figure S4C). A positive control sgRNA (ALU_112a) that was non-selective (Figure 3A) generated >99% growth inhibition in both cell lines (data not shown). Altogether, our results demonstrated that 4-9 sgRNAs, designed using our PAM discovery approach, were selectively toxic against targeted cells and produced significant cell death.

### Absence of off-target activity by patient-specific sgRNAs

We selected 7 of the 13 targets that we identified in Panc480 using our PAM discovery approach, confirmed functional targeting of individual sgRNAs, and cloned the corresponding sgRNAs into a multiplex sgRNA expression vector that expressed 7 sgRNAs simultaneously (designated as Panc480-MT7; figure S5A; table S10). Cells were harvested for deep sequencing at the targeted loci 14 days after transduction of Panc480-MT7. We detected cutting activity of all 7 sgRNAs in Panc480 Cas9-expressing cells, but not in its controls (Panc480 parental and Panc1002 Cas9-expressing cells) and corresponding normal (lymphoblasts) cells from the patient (Panc480-N; Figure S5B). To investigate potential off-target activity, we performed WGS on DNA extracted from Panc480 and Panc1002 Cas9-expressing cells 14 days post transduction (T14). Using two different programs, Cas-OFFinder (33) and IDT gRNA checker (34), we generated a list of potential sgRNA off-target sites of Panc480-MT7 and examined them on the WGS data. We found no evidence of Cas9 activity at these off-target sites (data not shown).

As an additional assessment of potential off-target activity, we used a somatic variant caller, MuTect2 (35), to identify novel indels present at 14 days and performed sequence alignment and homology comparison with the sgRNA sequences (36). We found that the indels novel to T14 did not exhibit significant homology to any of the 7 sgRNAs in Panc480-MT7 (table S11). The lowest possible mismatches were 4bp from the chr8:29032916 sgRNA and 5bp from the chr3:59525282 sgRNA (table S11), each with one occurrence only, and were unlikely to be targeted by the sgRNA as they were located at trinucleotide and dinucleotide repeat regions, respectively (data not shown). Novel indels at non-repetitive regions, with two examples included (figure S5C-D), were also unlikely to be targeted by Panc480-MT7 due to either absence of PAM, the distance between the potential sgRNA binding site and the indel, and/or the high number of mismatches, which have not been reported for Cas9 activity (15,17,37). These indels, present at low VAF, likely represent sequencing errors at repetitive regions, background heterogeneity in a bulk cell population, or ongoing genomic instability. Nonetheless, our results showed that these sgRNAs were highly specific to the intended targets in the targeted cell line and had no obvious off-target activity.

## Discussion

We present an efficient, cancer-specific PAM discovery approach that selectively kills cancer cells. We discovered that PCs, which generally have low mutational burden, contained >400 somatic PAMs as candidates for CRISPR-Cas9 targeting, significantly expanding the repertoire of targetable mutations in a given solid tumor. Since single-base mutations increase as a function of age (1,27), one could hypothesize that adult solid tumors, in general, contained hundreds of novel PAMs for subsequent selection of sgRNAs and targeting. This was supported by our mutational signature analyses which revealed aging signatures in most tumors and additional tissue-dependent factors, likely environmental, increasing the number of somatic PAMs. While it is conceivable that pediatric tumors might not contain as many somatic PAMs as adult cancers, we found that significant toxicity could be achieved with <10 sgRNAs, providing evidence that only a few sgRNAs would be needed to achieve selective killing. Optimization of sgRNA selection to maximize toxicity and strategies to enhance the inhibitory effect would be essential to broaden the applicability of this approach, especially in pediatric patients. For example, incorporating a DSB repair inhibitor increased the toxicity of the sgRNAs treated (figure S6). Combining our approach with existing therapies, such as immunotherapy and chemotherapy, should also be considered for synergistic effect.

As our strategy exploits the vast number of novel PAMs located in noncoding regions, it requires WGS analyses of both tumor and normal genomes. Most published data only includes exome sequencing, and obtaining corresponding normal for each tumor sample is still not common. WGS analyses are more computationally demanding than for WES, but the exponential decrease in sequencing costs (38) and enhancement of computing power have made these issues less concerning (39). As very few sgRNAs were needed to produce significant toxicity, one could potentially discover sufficient targetable PAMs from standard clinical workflow involving solid tumor panel sequencing. Generating cell lines from primary tumor samples and corresponding normal tissues for sgRNA screening continues to be challenging; however, this process has become much more routine with the advent of organoids. We have generated many cell lines from tumors and normal tissues of PCs, providing us with unique opportunities to study them extensively *in vitro* (23,40–42). This would also be particularly helpful for tumors with low cellularity (such as PCs (43)). Delivery of CRISPR-Cas therapeutics continues to be an active work of progress, with *in vivo* efficacy already demonstrated in the clinical setting and more delivery strategies under rapid development (44,45).

Off-target activity of CRISPR-Cas9 has been a general concern (46–49), with tolerance of up to 5 mismatches reported in the literature (15). We employed rigorous approaches to address this concern by performing WGS to examine for off-target activity of the sgRNAs under long term expression (14 days). We inspected potential off-target sites generated by two different sgRNA checkers, complemented with pairwise alignments on mutations identified by a somatic variant caller against the sgRNA sequences used. Although we didn’t detect any off-target activity, we recognized that the level of detection using WGS would not be as sensitive as targeted deep sequencing. As activity at one-mismatch sites was more commonly reported in the literature compared to 2-5 mismatches (16,17), we intentionally selected cell line-specific sgRNAs that had no 1-mismatch sites in the human genome for our co-culture assays. Meanwhile, strategies to mitigate off-target effects continue to evolve, with various approaches (such as improved sgRNA design, enhanced nuclease fidelity, and limiting exposure time of targeted cells to CRISPR-Cas9) have been proposed (16,49–51).

PAM-finding approaches have been published in a few studies (52,53). However, our approach is cancer- and, more importantly, patient-specific. This strategy presents a unique opportunity as a new precision medicine-based therapeutic tool that possesses the specificity of a targeted therapy, but without the restriction of a targetable protein and the drug development to target it (34). As cancer is a clonal disease, the distinct set of mutations found in the cancer initiating cell should be present in all primary tumor and metastatic sites, thus making genetic targeting of somatic PAMs a viable option for patients with multiple metastases (54). Since mutations are a universal feature of cancer, we envision applicability of our approach to a broad range of cancers.

## Acknowledgements

We thank the generosity of the patients who consented to providing their samples. We also acknowledge Drs. Ying Zou, Leslie Cope, Chien-Fu Hung, Elizabeth Jaffee, Lei Zheng, Feyruz V. Rassool, Richard Burkhart, Christine Iacobuzio-Donahue, Jessica Gucwa, Qingfeng Zhu, Shiwen Peng, and Toni Seppulla for helpful discussions. We thank Aparna Pallavajjala, Ada Tam, Raluca Yonescu, Laure Morsberger, Emily Adams, Lisa Haley, Anuj Gupta, Lee Blosser, Nicholas Ionta, Suping Chen, Kai Calder, Jormoh Bellepu, Jordyn Winters, and Nori Leybengrub for outstanding technical assistance. We also thank the ICGC-DACO investigators, particularly Drs. Anthony Gill, Nicola Waddell, John Pearson, Steven Gallinger, See Hoon Lee, and Rebecca Fitzgerald, for generating and sharing their data.

## Methods

### WGS-based PAM discovery and sgRNA design

Genomic DNA from tumors and corresponding normal tissues of Panc480, Panc504, and Panc1002 were whole genome sequenced and FASTQ files were aligned to hg38 using bwa v0.7.7 (mem, https://github.com/lh3/bwa) (55) to create BAM files. Picard-tools1.119 (RRID:SCR_006525) was used to add read groups as well as to remove duplicate reads. GATK v3.6.0 (RRID:SCR_001876, (56)) base call recalibration steps were used to create a final alignment file. MuTect2 v3.6.0 (56) was used to call somatic variants between the tumor-normal pairs. The default parameters and SnpEff (v4.1) (57) were used to annotate the passed variant calls.

PAMfinder (perl) was written to process VCFs based on their genome builds (hg38) to identify somatic variants that produced novel PAMs. Tumor (arrayT) and normal (arrayN) were specified based on column number, read depth was set at 18X (58), and VAF cutoff was modified based on the tumor purity (30% cutoff for 100% tumor purity). The 5’ and 3’ genomic sequences flanking the somatic variants were obtained from the FASTA of individual chromosomes to inspect whether novel Cs were adjacent to an existing C or novel Gs were adjacent to an existing G. The output contained information about the somatic variant, the potential sgRNA sequence along with the novel PAM, and specified whether the novel PAM was located on the plus or minus strand of the genome. Script is available on https://github.com/selinateh/PAMfinder.

Somatic mutations with VAF>95% were chosen to put through CRISPOR (59). Mutations that produced sgRNAs with >50 specificity score in CRISPOR were subsequently validated by PCR and Sanger sequencing (Primers Table 1).

### Somatic PAM discovery on ICGC samples

VCFs containing raw SNV calls generated via the GATK Mutect2 variant calling workflow were downloaded from the ICGC-ARGO Data Portal (60). These VCFs were sourced from four projects: APGI-AU (Australian Pancreatic Cancer Genome Initiative; N=44), LUCA-KR (Personalised Genomic Characterisation of Korean Lung Cancers; N=29), PACA-CA (Pancreatic Cancer Harmonized “Omics” analysis for Personalized Treatment; N=130), and OCCAMS-GB (Oesophageal Cancer Clinical and Molecular Stratification; N=388). To correct for tumor purity, we used the VCF for each sample, comprising variants for tumor and normal tissue, as input to the R package FACETS (61) to calculate tumor purity. Then, we used both the VCF and the tumor purity data as input to PAMfinder to identify base substitutions that produced novel PAMs. The VAF cutoff for somatic PAM calling was % tumor purity x 30% (e.g. if the tumor purity was 50%, the VAF cutoff for somatic PAM calling was 15%). Finally, % novel PAM was calculated by dividing the number of novel PAM by the total number of base substitutions.

### Computing mutational profiles and COSMIC signature contribution

MutationalPatterns R package was used to calculate SBS mutational profiles for each VCF file and to compute the contribution of each of 60 COSMIC signatures to these profiles (24,62). The 12 displayed COSMIC signatures represented the 12 most frequently appearing amongst the top five signatures across all samples, when signatures were ordered from the greatest contribution to the least contribution within each sample.

### Multitarget sgRNA design and cell viability assay

Chromosome range was entered into CRISPOR (59) 2kb at a time starting at chr1:0-2000 and ending at chr1:100,248,000-100,250,000 based on hg38. sgRNAs that had different number of perfect target sites were selected from the pool of sgRNA options generated by CRISPOR based on the following criteria: (1) none of the perfect target sites and potential off-target sites target exons; (2) Doench’16 efficiency score >50%, and (3) the number of off-targets that have no mismatches in the 12bp adjacent to the PAM (SEED region) is <10 (16). Sequences of non-targeting control sgRNAs were obtained from Doench *et al.* (NT) and Chiou *et al.* (NT2) (16,63). Positive control sgRNAs were designed by inserting LINE-1 and Alu element sequences to CRISPOR.

Cas9-expressing PC cells were transduced with lentivirus at MOI10. The cells were split into 96-well plates in 1:1000 dilution for clonogenicity. When cells in non-targeting controls reached full confluence (1-2 months), alamarBlue Cell Viability Reagent (ThermoFisher) was added and BMG POLARstar Optima microplate reader was used for fluorescence reading to assess cell viability. Excitation was set at 544nm and emission at 590nm, with a gain of 1000 and required value of 90%.

### Co-culture assays

Parental cells and/or cells that expressed either mApple or mNeon-Green fluorophores were co-cultured at different ratios under antibiotic selection. Puromycin selection was for 7 days at 1ug/mL and hygromycin selection was for 14 days at 200ug/mL. Proportions of mApple-expressing cells (Ex/Em: 561/620 nm) and/or mNeon-Green-expressing cells (Ex/Em: 488/530 nm) post-transduction of sgRNAs were measured and analyzed at different time points using Attune NxT Flow Cytometer (ThermoFisher) and FCS Express 7 (De Novo Software).

### Mouse-human NGS assay

The *RC3H2* gene was selected as the mouse and human orthologs as they differ by a 3bp indel followed by 3 SBSs (figure S3C). Primers for unbiased PCR amplification of the locus in mouse and human DNA were previously developed by Lin *et. al*., designated as primer pair 45 (64). For this assay, a 101bp amplicon in the *RC3H2* gene was amplified with primers containing Illumina adaptor sequences (Primers Table 2). Amplicons were subjected to NGS (see Supplementary Methods), and FASTQ files were aligned to the hg19 genome using bwa 0.7.17(65) and visualized in IGV. Human and mouse reads were quantified since the 3bp-shorter mouse sequence maps as a deletion in the human genome. For assay validation, mouse DNA and human DNA were mixed at varying ratios and assayed (figure S3D).

### Functional testing of multiplex sgRNA expression plasmids

The targeted cell line and non-targeted cell line(s) were transduced at MOI 10 with lentivirus expressing the multiplex vectors. 14-21 days after transduction, cells were harvested and genomic DNA extracted using QIAmp UCP DNA Micro kit (QIAGEN). The targeted loci were PCR amplified with NGS adaptors and sent for amplicon sequencing (Primers Table 1 “Panc480 mutation validation” & 3, see Supplementary Methods). The sequencing data was analyzed for the percent of edited reads by CRISPResso2 (66).

### WGS analyses of potential off-target sites

Two replicates of Panc480 Cas9-expressing cell pellets and one replicate of Panc1002 Cas9-expressing cell pellet from the Panc480-MT7 functional testing assay were used for this analysis. Cells were harvested 14 days after transduction (T14) or the day of transduction (T0). DNA was whole genome sequenced and FASTQ files were aligned to hg19 using bwa v0.7.7 (55) to create BAM files. Picard-tools1.119 (RRID:SCR_006525) was used to add read groups as well as to remove duplicate reads. GATK v3.6.0 (RRID:SCR_001876, (56)) base call recalibration steps were used to create a final alignment file.

sgRNA sequences were put through Cas-OFFinder (33) to identify potential off-target sites including ones with non-canonical NAG PAM and ones with 1-4 mismatches. sgRNA sequences were also uploaded to IDT CRISPR-Cas9 gRNA checker (34) to obtain a second list of potential off-target sites. Then, we examined each site on IGV for Cas9-induced mutation signatures.

MuTect2 v3.6.0 (56) was used to call somatic variants between the T14-T0 pairs. The default parameters and SnpEff (v4.1) (57) were used to annotate the passed variant calls. From the list of variants generated, we performed homology analysis with an R script that performed the following steps: 1) Read in an Excel file containing one mutation per row. 2) Obtain the forward and reverse strand sequences from the hg19 genome between the start – 50 bp and stop + 50 bp positions of the locus. 3) Align each locus’s forward and reverse sequences to the target sgRNA with no gaps using the Smith-Waterman algorithm. 4) Determine the number of mismatches between the sgRNA and the nearest matching piece of DNA within each junctions. 5) Output the original information along with new columns displaying the mismatches between each junction and the sgRNA into a new Excel file. EMBOSS Needle was used to illustrate the alignments between the sgRNA sequences and the potential target sequences with the lowest number of mismatches (67).

### Statistical analysis

The appropriate statistical tests were performed in GraphPad Prism (Version 9.2.0, RRID:SCR_002798) and stated in the legends of figures. For all statistically significant results, * indicates *P<0.05*, ** indicates *P<0.01*, *** indicates *P<0.001*, and **** indicates *P<0.0001*.

## Data availability

The authors confirm that the data supporting the findings of this study are available within the article and its supplementary materials when applicable. ICGC data that support the findings of this study are available in the Cancer Genome Collaboratory at https://cancercollaboratory.org/. These data were derived from the ICGC ARGO platform. Plasmids constructed had been deposited at Addgene. Plasmids expressing various specific sgRNAs are available upon request from the corresponding author.

