## Supplementary material for "CRISPR-Cas9 for selective targeting of somatic mutations in pancreatic cancers": figure S: Primers Table 1-7.docx

Table 1. Primers for PCR and Sanger validation of novel base substitutions discovered from WGS approach.

Table 2. Primers used for mouse-human NGS assay.

Table 3. Primers used for PCR and NGS of Panc10.05 quad targets.

Table 4. Primers for PCR and Sanger validation of novel SVs.

Table 5. Primers for Cas9-mApple, Cas9-mNeonGreen, and dCas9-EGFP plasmid constructions and validations.

Table 6. Primers for Cas9 activity assay.

Table 7. Primers involved in multiplex sgRNA vector construction.

Table 1**. Primers for PCR and Sanger validation of novel base substitutions discovered from WGS approach.**

| **Primer name** | **Purpose** | **Sequence** |
| --- | --- | --- |
| Panc480_chr3:537601_Fwd | Panc480 mutation validation | TGAGACTGTATTTGTGGGCCA |
| Panc480_chr3:59525282_Fwd | Panc480 mutation validation | GGCCCTCACCATGTAAAAGG |
| Panc480_chr18:1819017_Fwd | Panc480 mutation validation | ACTGGGAAGTTGGGTCTTCA |
| Panc480_chrX:3982448_Fwd | Panc480 mutation validation | TGGAGGTAGGATATTACAGGGAA |
| Panc480_chr19:58564841_Fwd | Panc480 mutation validation | GCCATCCACTCACTACAGGT |
| Panc480_chr8:29032916_Fwd | Panc480 mutation validation | TGGAAGGCTAGAGGAAGCTG |
| Panc480_chr6:124767224_Fwd | Panc480 mutation validation | TGTGTGCCTTCAAAATGGGG |
| Panc480_chr6:55808003_Fwd | Panc480 mutation validation | TGAAGCATACATTCTGGAGGTT |
| Panc480_chr11:64364029_Fwd | Panc480 mutation validation | TGGATGAACTGGATGGATGA |
| Panc480_chr6:92757856_Fwd | Panc480 mutation validation | TGCCTAGTCCAGTAATGCGA |
| Panc480_chr17:5377742_Fwd | Panc480 mutation validation | ACACCATGGCCTCATCTATCA |
| Panc480_chr4:131074842_Fwd | Panc480 mutation validation | TGCTCTCAACTTTCCCTGGA |
| Panc480_chr8:201457_Fwd | Panc480 mutation validation | GGGGGATGGTCATGAGATTT |
| Panc480_chr3:86665957_Fwd | Panc480 mutation validation | CCTGCCCCAGTGAAATCAGT |
| Panc480_chr9:15347394_Fwd | Panc480 mutation validation | AGGCAGCTAGAGTTCACAGG |
| Panc480_chr9:110569399_Fwd | Panc480 mutation validation | GCAGAGGGGAGCTCTTTTCT |
| Panc480_chr1:34085551_Fwd | Panc480 mutation validation | CCATTCCTCTCCACACTCCA |
| Panc480_chr3:537601_rev | Panc480 mutation validation | AGCACGCAATATTACTGGGAAC |
| Panc480_chr3:59525282_rev | Panc480 mutation validation | TGACCACCACATCCAGGAT |
| Panc480_chr18:1819017_rev | Panc480 mutation validation | CACTCCCAAGAACGCAGAAT |
| Panc480_chrX:3982448_rev | Panc480 mutation validation | ACCATCGTTTTAAAAGGTGCAA |
| Panc480_chr19:58564841_rev | Panc480 mutation validation | GCTCGAGATCACAGTCCCTT |
| Panc480_chr8:29032916_rev | Panc480 mutation validation | ATGTGCGGTGGTAGGAGAAG |
| Panc480_chr6:124767224_rev | Panc480 mutation validation | AGCAATATGGAGGAACAAAAGCA |
| Panc480_chr6:55808003_rev | Panc480 mutation validation | GTCATCCACTTCATCCACTTCA |
| Panc480_chr11:64364029_rev | Panc480 mutation validation | AGGAGTGGCTGCAAATTGTT |
| Panc480_chr6:92757856_rev | Panc480 mutation validation | CGGTATAGTTTCCACAGCAGG |
| Panc480_chr17:5377742_rev | Panc480 mutation validation | CAGTTTGCCAGTGGTTCCTC |
| Panc480_chr4:131074842_rev | Panc480 mutation validation | CACCGAGTTTGAGATGCCTG |
| Panc480_chr8:201457_rev | Panc480 mutation validation | TGATCCAGTGTGGGTGAGAA |
| Panc480_chr3:86665957_rev | Panc480 mutation validation | GGAGAGTGTACCCTGTTGCT |
| Panc480_chr9:15347394_rev | Panc480 mutation validation | GCCCCGCTACTGAGAGAATA |
| Panc480_chr9:110569399_rev | Panc480 mutation validation | ACCTCATCTCCCTGCTATGC |
| Panc480_chr1:34085551_rev | Panc480 mutation validation | TCAGCCTCATCTTTCTCCCA |
| Panc1002_chr3:41255526_fwd | Panc1002 mutation validation | ACTTGACATGTATGGTGGGG |
| Panc1002_chr3:76569799_fwd | Panc1002 mutation validation | GGATTTTACAGCTGGAAGGGATC |
| Panc1002_chr4:32408343_fwd | Panc1002 mutation validation | GCAACATTGCATGTTCAGAAA |
| Panc1002_chr4:117677347_fwd | Panc1002 mutation validation | CGGTAGCTTGGATGACAGAA |
| Panc1002_chr4:180416652_fwd | Panc1002 mutation validation | GGCCCTACCCATACCTACTG |
| Panc1002_chr4:180746369_fwd | Panc1002 mutation validation | TAGGACTACAGCAGCACACC |
| Panc1002_chr6:123690025_fwd | Panc1002 mutation validation | TCCATTCCTTGTTCTTGCCAC |
| Panc1002_chr6:153579209_fwd | Panc1002 mutation validation | CCAAGCAACATAAAGCAGCA |
| Panc1002_chrX:28266415_fwd | Panc1002 mutation validation | TCTTTCTCCTAGATCTGGACACT |
| Panc1002_chrX:56623848_fwd | Panc1002 mutation validation | GCTGCCTTTCTTCCAGTGAT |
| Panc1002_chrX:116828813_fwd | Panc1002 mutation validation | AGGCTCCACTGCTTCTGTGT |
| Panc1002_chr8:12552195_fwd | Panc1002 mutation validation | TCCTGGGGCAATTTTACTTTT |
| Panc1002_chr8:47456593_fwd | Panc1002 mutation validation | GCTCACCCACTTTCCATTCA |
| Panc1002_chr8:81741154_fwd | Panc1002 mutation validation | TCTGCCCCAACATGAGACTT |
| Panc1002_chr9:23649543_fwd | Panc1002 mutation validation | TGTCCACACCTACAATCCTGA |
| Panc1002_chr11:55366717_fwd | Panc1002 mutation validation | TCAGTTGTTTCACAGATCTGCA |
| Panc1002_chr12:47771504_fwd | Panc1002 mutation validation | GTGCAGCTTCACTCCTCACA |
| Panc1002_chr18:58907286_fwd | Panc1002 mutation validation | CAATTGCAACGGGAATTCTT |
| Panc1002_chrY:17028622_fwd | Panc1002 mutation validation | GCAGATAATGACCTTCCTATTGC |
| Panc1002_chr3:15793085_fwd | Panc1002 mutation validation | GGTAGAGAAAAGCCCTGAGGA |
| Panc1002_chr3:27365096_fwd | Panc1002 mutation validation | GAGAACGGGAGGATTCTGG |
| Panc1002_chr4:45316432_fwd | Panc1002 mutation validation | TGCATCACAAGGGTTATTGC |
| Panc1002_chr4:58746119_fwd | Panc1002 mutation validation | ATGCAACCTTTTGTGTTCCA |
| Panc1002_chr4:63298774_fwd | Panc1002 mutation validation | TGTGGCACAGATTTATTAGCAGA |
| Panc1002_chr7:158427297_fwd | Panc1002 mutation validation | ACAGGCACAACCATCCATTT |
| Panc1002_chrX:9204373_fwd | Panc1002 mutation validation | ATGCCTGCATTTACCACCAT |
| Panc1002_chrX:99446566_fwd | Panc1002 mutation validation | CCAATTTTAGGCATGCAGGT |
| Panc1002_chr8:88685752_fwd | Panc1002 mutation validation | GGCAAATGTTCCCTGATGTT |
| Panc1002_chr9:15744747_fwd | Panc1002 mutation validation | GCCAATCATGTGCCTCTCTT |
| Panc1002_chr17:876863_fwd | Panc1002 mutation validation | TTTCCCAGGCTTCGTCGAT |
| Panc1002_chr18:39354909_fwd | Panc1002 mutation validation | GCGGGGATTTGCACAGAATT |
| Panc1002_chr18:51635625_fwd | Panc1002 mutation validation | GCACTCGAAGGCTTCTCC |
| Panc1002_chr19:5559720_fwd | Panc1002 mutation validation | TCAATCAAGTGAGACAGGGCT |
| Panc1002_chr21:24912568_fwd | Panc1002 mutation validation | CATGGGAGGCTGGATTCATT |
| Panc1002_chr3:41255526_rev | Panc1002 mutation validation | CTCCCCATAGCTAAGGACCA |
| Panc1002_chr3:76569799_rev | Panc1002 mutation validation | GTCAAGATGTGGACTACTAGCA |
| Panc1002_chr4:32408343_rev | Panc1002 mutation validation | GCCAAATCGGAAACAAAGAA |
| Panc1002_chr4:117677347_rev | Panc1002 mutation validation | CAATGTAAGTGGGCAGCAGA |
| Panc1002_chr4:180416652_rev | Panc1002 mutation validation | ACCAAGGCTAAAGATCAGTGAT |
| Panc1002_chr4:180746369_rev | Panc1002 mutation validation | TCATTGGTATTTGGAGCTTTGC |
| Panc1002_chr6:123690025_rev | Panc1002 mutation validation | CCAGCCTCTAGAACTGTGGA |
| Panc1002_chr6:153579209_rev | Panc1002 mutation validation | ATGGTGTGTCAGACGCTGTT |
| Panc1002_chrX:28266415_rev | Panc1002 mutation validation | GGTAAATAACTTTGTCCTGGGTG |
| Panc1002_chrX:56623848_rev | Panc1002 mutation validation | GAAATTCTTCCTGCCAGCAC |
| Panc1002_chrX:116828813_rev | Panc1002 mutation validation | TGGTGGTGTTGGTGATTCAG |
| Panc1002_chr8:12552195_rev | Panc1002 mutation validation | TGGTGGTGTTGGTGATTCAG |
| Panc1002_chr8:47456593_rev | Panc1002 mutation validation | TGCTTGCTTAAACTCCTCAGT |
| Panc1002_chr8:81741154_rev | Panc1002 mutation validation | GGGTGACAATCTTCCTGTGG |
| Panc1002_chr9:23649543_rev | Panc1002 mutation validation | GTTCCTTCAATTGCCGATGT |
| Panc1002_chr11:55366717_rev | Panc1002 mutation validation | CAGCTCATCCAGAACCCAGA |
| Panc1002_chr12:47771504_rev | Panc1002 mutation validation | ATGCTGCTGTGATCGTTTTG |
| Panc1002_chr18:58907286_rev | Panc1002 mutation validation | GGAAAGTGGTGTCCAGGATG |
| Panc1002_chrY:17028622_rev | Panc1002 mutation validation | CATGAATTACAAGGGCAGCAA |
| Panc1002_chr3:15793085_rev | Panc1002 mutation validation | ATAGGCGTACCCCTGAATCC |
| Panc1002_chr3:27365096_rev | Panc1002 mutation validation | AAAGACCTTTGAAGGATGCAA |
| Panc1002_chr4:45316432_rev | Panc1002 mutation validation | TGGATTCCAGAAATTGTTTTTGA |
| Panc1002_chr4:58746119_rev | Panc1002 mutation validation | GCTATTCATTAGCGGGGACA |
| Panc1002_chr4:63298774_rev | Panc1002 mutation validation | AAAGGCTTAGTGCTGACCTTACA |
| Panc1002_chr7:158427297_rev | Panc1002 mutation validation | CATGGGCAGTTTGCTTTACC |
| Panc1002_chrX:9204373_rev | Panc1002 mutation validation | TTTCCAAGGTGATGACCACA |
| Panc1002_chrX:99446566_rev | Panc1002 mutation validation | AGAAGGCCCTTTCATCATCA |
| Panc1002_chr8:88685752_rev | Panc1002 mutation validation | AACTGGATTGGTTGCTGCTT |
| Panc1002_chr9:15744747_rev | Panc1002 mutation validation | ACACTGTATTTCGCTTACATGCA |
| Panc1002_chr17:876863_rev | Panc1002 mutation validation | TGGGTGACAGAGCAAGACT |
| Panc1002_chr18:39354909_rev | Panc1002 mutation validation | GGCTCCTCCTCCCTACAAAT |
| Panc1002_chr18:51635625_rev | Panc1002 mutation validation | TCATCCCTTTGTCCAGCAGA |
| Panc1002_chr19:5559720_rev | Panc1002 mutation validation | TGTCCTCATTTCCCTGTGCA |
| Panc1002_chr21:24912568_rev | Panc1002 mutation validation | AGACACGTAACGGCAGATGT |
| Panc504_chr1:90925384_fwd | Panc504 mutation validation | TCTTTGTCTTGTGCATGGCG |
| Panc504_chr1:109094826_fwd | Panc504 mutation validation | CTTAGAAAAGGCACAGCATAGG |
| Panc504_chr4:96761136_fwd | Panc504 mutation validation | GCTCCAGGGTTTAACAGGGA |
| Panc504_chr4:147513098_fwd | Panc504 mutation validation | GCCAGCCTTGAAGTGTGTC |
| Panc504_chrX:10649926_fwd | Panc504 mutation validation | GCACATCCAAATTTATTCACACG |
| Panc504_chrX:137303674_fwd | Panc504 mutation validation | GAACAACACCAGGCACATAGT |
| Panc504_chrX:141322626_fwd | Panc504 mutation validation | GGAATTCCTGACTCCAAAACA |
| Panc504_chr9:10209960_fwd | Panc504 mutation validation | CTGGTGCTTTTGTTTTGATTAGG |
| Panc504_chr9:77440886_fwd | Panc504 mutation validation | AGGCAACAGGACATTTCAGG |
| Panc504_chr9:105373293_fwd | Panc504 mutation validation | GCTGTTCCAATACAAGCCCC |
| Panc504_chr9:133876782_fwd | Panc504 mutation validation | TCTGGTCCCATAACTGCACA |
| Panc504_chr10:4171262_fwd | Panc504 mutation validation | TCTGGAGAACAAAGGCATTCC |
| Panc504_chr13:107175748_fwd | Panc504 mutation validation | GGTTCCTGACTTCCATACGG |
| Panc504_chr18:39014688_fwd | Panc504 mutation validation | GGGAGGGAGGGAAGAAACAA |
| Panc504_chr18:48358086_fwd | Panc504 mutation validation | TGCATTTCTTATTTCCCAGCAAC |
| Panc504_chr18:63239834_fwd | Panc504 mutation validation | AGCTGTGCAGGATTGAATTCT |
| Panc504_chr21:23671417_fwd | Panc504 mutation validation | ATGACCAAAATGAGAAATTATTAGC |
| Panc504_chr1:25383677_fwd | Panc504 mutation validation | GTATGCCAGGAGCCAGGTT |
| Panc504_chr1:30192392_fwd | Panc504 mutation validation | CTTGGGTATGTGCCTTGCTC |
| Panc504_chr1:73167766_fwd | Panc504 mutation validation | GCATGTGTTTACCTGGCCTAC |
| Panc504_chr1:82861966_fwd | Panc504 mutation validation | CCTAAGGGTGTGACTCCAGA |
| Panc504_chr4:32481045_fwd | Panc504 mutation validation | CATCACGCCCGGCTAATTTT |
| Panc504_chr4:98124868_fwd | Panc504 mutation validation | GAGCTTTTGAATGGTGACTGGA |
| Panc504_chr4:146038680_fwd | Panc504 mutation validation | CAAGCGCCTATGGAGTTGTC |
| Panc504_chr4:177915089_fwd | Panc504 mutation validation | AGAAACCAGTGAAGGATCTCC |
| Panc504_chr4:189873183_fwd | Panc504 mutation validation | GGGCAATAAACATGAAAAGTGGT |
| Panc504_chr5:50335067_fwd | Panc504 mutation validation | ACAGCCCCAATCTGTTTCAC |
| Panc504_chr5:76384387_fwd | Panc504 mutation validation | TAGAGGAGTTGGGGGAAGGT |
| Panc504_chr5:117548593_fwd | Panc504 mutation validation | TCATCCCGAGAGTTATATCCCC |
| Panc504_chr7:97304833_fwd | Panc504 mutation validation | AAGATCAAGCCAGCCACAAT |
| Panc504_chr7:110208712_fwd | Panc504 mutation validation | CATCAACTCACTCACAGGCAG |
| Panc504_chr7:137081417_fwd | Panc504 mutation validation | GATGTGCTGGCATGTGGAC |
| Panc504_chrX:19715766_fwd | Panc504 mutation validation | GCTGCGGGACATAGAACTGT |
| Panc504_chrX:22650252_fwd | Panc504 mutation validation | TGACCCTGGAATTCACCTGC |
| Panc504_chrX:27834613_fwd | Panc504 mutation validation | TGTATCTGCGCCAAGGGAAA |
| Panc504_chrX:105633682_fwd | Panc504 mutation validation | TTTTGAGTGAACGTGGCAGC |
| Panc504_chrX:113360530_fwd | Panc504 mutation validation | AGGATTACTGATTGGGCCACT |
| Panc504_chr8:15708017_fwd | Panc504 mutation validation | AGGTTTGTTCTCCCATAGTTGA |
| Panc504_chr9:128664573_fwd | Panc504 mutation validation | AGATGTTTGCTCCAAGAACCT |
| Panc504_chr13:67584092_fwd | Panc504 mutation validation | ACAAAGACATGCAACAGATCACA |
| Panc504_chr13:70467817_fwd | Panc504 mutation validation | AGCAAACAAAAGAACCACTAGCT |
| Panc504_chr13:92785652_fwd | Panc504 mutation validation | AGGGTGTCGTACTAAATGGGA |
| Panc504_chr18:69135730_fwd | Panc504 mutation validation | CCAAGGTTAGGTGTGGGGAA |
| Panc504_chr22:34609948_fwd | Panc504 mutation validation | GCTAAGGTGATCAACAAGTTTCC |
| Panc504_chr21:29359027_fwd | Panc504 mutation validation | AGATCTCCCTTTTGTTGGTTGA |
| Panc504_chr1:90925384_rev | Panc504 mutation validation | CAGGGATGTGTGGGAGATGA |
| Panc504_chr1:109094826_rev | Panc504 mutation validation | GGTACGCACTCAATAGCTGG |
| Panc504_chr4:96761136_rev | Panc504 mutation validation | GGGTGATAGAGGCAGGTCC |
| Panc504_chr4:147513098_rev | Panc504 mutation validation | CCTTTACCCTCAAGTGCTTTCC |
| Panc504_chrX:10649926_rev | Panc504 mutation validation | TGAGTGTCTATTAAGTGCCAGTG |
| Panc504_chrX:137303674_rev | Panc504 mutation validation | CAGACCACCTATGACTAGAGCA |
| Panc504_chrX:141322626_rev | Panc504 mutation validation | GTCCCCCTTCCTCAATCAAT |
| Panc504_chr9:10209960_rev | Panc504 mutation validation | TGTTTTCAGAAATAAACTTTTTCACC |
| Panc504_chr9:77440886_rev | Panc504 mutation validation | CTCTGGGAATTGTGGTCGTT |
| Panc504_chr9:105373293_rev | Panc504 mutation validation | GGTGCTACTTGTCTCTCAGC |
| Panc504_chr9:133876782_rev | Panc504 mutation validation | CATGAAATGGGAACGGTAGG |
| Panc504_chr10:4171262_rev | Panc504 mutation validation | CCACAGACAGAGTAGGACAGA |
| Panc504_chr13:107175748_rev | Panc504 mutation validation | CAGCACATCCTCCTTCCTCC |
| Panc504_chr18:39014688_rev | Panc504 mutation validation | TCCCACCGTTCTCTGATCAT |
| Panc504_chr18:48358086_rev | Panc504 mutation validation | AGTTGCTGTGGAGACCTTCA |
| Panc504_chr18:63239834_rev | Panc504 mutation validation | ACTTGTTTCATGCCCTTGTTTT |
| Panc504_chr21:23671417_rev | Panc504 mutation validation | TTGGTTGTGCTTCTTGTTGAA |
| Panc504_chr1:25383677_rev | Panc504 mutation validation | TCGAGAAGGGAAAGATTGGA |
| Panc504_chr1:30192392_rev | Panc504 mutation validation | TGGTGATGGAGGCAATGACT |
| Panc504_chr1:73167766_rev | Panc504 mutation validation | ATAGGAGGGAGGCACAAGTG |
| Panc504_chr1:82861966_rev | Panc504 mutation validation | GGTGATAAAGCGACCTTGAGT |
| Panc504_chr4:32481045_rev | Panc504 mutation validation | GTACAGAGTCTCGGATGCTTTT |
| Panc504_chr4:98124868_rev | Panc504 mutation validation | CACACCACTCCATTTGTCTGT |
| Panc504_chr4:146038680_rev | Panc504 mutation validation | TGCTCAGTGATTAAATTCCAAGG |
| Panc504_chr4:177915089_rev | Panc504 mutation validation | ATGCTATCATCATGGGCCCC |
| Panc504_chr4:189873183_rev | Panc504 mutation validation | TGGACAGACATTTGGGGTGA |
| Panc504_chr5:50335067_rev | Panc504 mutation validation | TCCAGGTGACTTGATGTAGCA |
| Panc504_chr5:76384387_rev | Panc504 mutation validation | CAGCAGCAAAAGATGAGCAG |
| Panc504_chr5:117548593_rev | Panc504 mutation validation | TCTGTCCTAATGCCCTTCCA |
| Panc504_chr7:97304833_rev | Panc504 mutation validation | AGCTCTGGAAGTAGGCATTGA |
| Panc504_chr7:110208712_rev | Panc504 mutation validation | CCACTGAGGGTATTGGGACA |
| Panc504_chr7:137081417_rev | Panc504 mutation validation | TGAGTTGGTGTGGAGAGGAA |
| Panc504_chrX:19715766_rev | Panc504 mutation validation | TAGCACCCCAGATCTCAGTG |
| Panc504_chrX:22650252_rev | Panc504 mutation validation | GATTGAACCCTCATCATTTGCC |
| Panc504_chrX:27834613_rev | Panc504 mutation validation | CCCCGCTGCACTCAATAAC |
| Panc504_chrX:105633682_rev | Panc504 mutation validation | GCATTCTCTCACTCAAGCACA |
| Panc504_chrX:113360530_rev | Panc504 mutation validation | TGGCTGTTCAGATATTGGATTCA |
| Panc504_chr8:15708017_rev | Panc504 mutation validation | GGGGAAAGAGATGAGAAGAGAGA |
| Panc504_chr9:128664573_rev | Panc504 mutation validation | AGAGTCATTGTCTACGATCCCA |
| Panc504_chr13:67584092_rev | Panc504 mutation validation | TGCTCTTCACATTTCCTGAACA |
| Panc504_chr13:70467817_rev | Panc504 mutation validation | GCCATTTCCAGAATTGAGACCA |
| Panc504_chr13:92785652_rev | Panc504 mutation validation | TGCCTCCTTGAATGAACTGTG |
| Panc504_chr18:69135730_rev | Panc504 mutation validation | AGAGAGAAACACTAGTAGCCTGA |
| Panc504_chr22:34609948_rev | Panc504 mutation validation | GCGTAACTGCTAGAAGAAGAGA |
| Panc504_chr21:29359027_rev | Panc504 mutation validation | AAGTCACTGGGAAGCAGTCA |

**Table 2. Primers used for mouse-human NGS assay.**

| **Primer name** | **Sequence** |
| --- | --- |
| NGS-RC3H2-45-Lib-Fwd-1 | AATGATACGGCGACCACCGAGATCTACACTCTTTCCCTACACGACGCTCTTCCGATCTTAAGTAGAGactaagtcaaggctactgtg |
| NGS-RC3H2-45-Lib-Fwd-2 | AATGATACGGCGACCACCGAGATCTACACTCTTTCCCTACACGACGCTCTTCCGATCTATCATGCTTAactaagtcaaggctactgtg |
| NGS-RC3H2-45-Lib-Fwd-3 | AATGATACGGCGACCACCGAGATCTACACTCTTTCCCTACACGACGCTCTTCCGATCTGATGCACATCTactaagtcaaggctactgtg |
| NGS-RC3H2-45-Lib-Fwd-4 | AATGATACGGCGACCACCGAGATCTACACTCTTTCCCTACACGACGCTCTTCCGATCTCGATTGCTCGACactaagtcaaggctactgtg |
| NGS-RC3H2-45-Lib-Fwd-5 | AATGATACGGCGACCACCGAGATCTACACTCTTTCCCTACACGACGCTCTTCCGATCTTCGATAGCAATTCactaagtcaaggctactgtg |
| NGS-RC3H2-45-Lib-KO-Rev- 1 | CAAGCAGAAGACGGCATACGAGATTC GCCTTGGTGACTGGAGTTCAGACGTGTGCTCTTCCGATCTttctggtgtcagtatggaag |
| NGS-RC3H2-45-Lib-KO-Rev- 2 | CAAGCAGAAGACGGCATACGAGATAT AGCGTCGTGACTGGAGTTCAGACGTGTGCTCTTCCGATCTttctggtgtcagtatggaag |
| NGS-RC3H2-45-Lib-KO-Rev- 3 | CAAGCAGAAGACGGCATACGAGATGA AGAAGTGTGACTGGAGTTCAGACGTGTGCTCTTCCGATCTttctggtgtcagtatggaag |
| NGS-RC3H2-45-Lib-KO-Rev- 4 | CAAGCAGAAGACGGCATACGAGATAT TCTAGGGTGACTGGAGTTCAGACGTGTGCTCTTCCGATCTttctggtgtcagtatggaag |
| NGS-RC3H2-45-Lib-KO-Rev- 5 | CAAGCAGAAGACGGCATACGAGATCG TTACCAGTGACTGGAGTTCAGACGTGTGCTCTTCCGATCTttctggtgtcagtatggaag |

**Table 3. Primers used for PCR and NGS of Panc10.05 quad targets.**

| **Name** | **Sequence** |
| --- | --- |
| NGS-Panc10.05-chr13:67159870-Fwd1 | AATGATACGGCGACCACCGAGATCTACACTCTTTCCCTACACGACGCTCTTCCGATCTGATGCACATCTCCTTAACGCATTCCCCAGTA |
| NGS-Panc10.05-chr13:67159870-Rev1 | CAAGCAGAAGACGGCATACGAGATGA AGAAGTGTGACTGGAGTTCAGACGTGTGCTCTTCCGATCTTTTGTTCACCTCTGCACACC |
| NGS-Panc10.05-chr13:98046824-Fwd1 | AATGATACGGCGACCACCGAGATCTACACTCTTTCCCTACACGACGCTCTTCCGATCTGATGCACATCTTGGGATCAATGGCATCAGTA |
| NGS-Panc10.05-chr13:98046824-Rev1 | CAAGCAGAAGACGGCATACGAGATGA AGAAGTGTGACTGGAGTTCAGACGTGTGCTCTTCCGATCTCCTCATCACCCTCTCTTTGC |
| NGS-Panc10.05-chr3:67534508-Fwd1 | AATGATACGGCGACCACCGAGATCTACACTCTTTCCCTACACGACGCTCTTCCGATCTGATGCACATCTGCTTGCCATGAATGGAGTTT |
| NGS-Panc10.05-chr3:67534508-Rev1 | CAAGCAGAAGACGGCATACGAGATGA AGAAGTGTGACTGGAGTTCAGACGTGTGCTCTTCCGATCTGAGAGTGGGATGAGGTTCCA |
| NGS-Panc10.05-chr3:76973238-Fwd1 | AATGATACGGCGACCACCGAGATCTACACTCTTTCCCTACACGACGCTCTTCCGATCTGATGCACATCTTGCTTAAGAATTTGAGCGACA |
| NGS-Panc10.05-chr3:76973238-Rev1 | CAAGCAGAAGACGGCATACGAGATGA AGAAGTGTGACTGGAGTTCAGACGTGTGCTCTTCCGATCTCGTGTTCTCATCACCTTCCA |

Table 4**. Primers for PCR and Sanger validation of novel SVs.**

| **Forward primer^*^** | **Sequence^#^** | **Reverse primer^*^** | | **Sequence^#^** |
| --- | --- | --- | --- | --- |
| PANC480_Chr1:174M_td Fwd | GTAAAACGACGGCCAGCTCTTTGGCTGATGTTCC | PANC480_Chr1:174M_td Rev | | CAGGAAACAGCTATGACTCTGCACATAACGGTGGA |
| PANC480_chr1_154d_st1_fwd | GTAAAACGACGGCCAGAAGAATCGCCTGAACCTGGG | PANC480_chr1_154d_st1_rev | | GCCTGTCCCTTGTTTCCTTG |
| PANC480_Chr1:222M_t Fwd | GTAAAACGACGGCCAGTCTCAAAGTTACACGTCA | PANC480_Chr1:222M_t Rev | | CAGGAAACAGCTATGACAGTAGAGAAGCTTGAAAT |
| PANC480_chr1_248t_st1_fwd | GTAAAACGACGGCCAGACTACCACTCCTTCATCCCC | PANC480_chr1_248t_st1_rev | | TGCACACATCACAAAGAAGTTTC |
| PANC480_chr2_26d_st1_fwd | GTAAAACGACGGCCAGGTTCACCATCTTAGCCACAGG | PANC480_chr2_26d_st1_rev | | CCCAGGCTGTTCTCGAAAAC |
| PANC480_Chr2:149M_D_FWD | GTAAAACGACGGCCAGAAAGAGTGTGACGGAGGG | PANC480_Chr2:149M_D_REV | | CAGGAAACAGCTATGACATGAAAACAGTGAAATAT |
| PANC480_chr2_221d_st1_fwd | GTAAAACGACGGCCAGTATTTGATGAGGGCCAGTGC | PANC480_chr2_221d_st1_rev | | GGAACCTCTGCTCTTCATGAC |
| PANC480_Chr2:164M_td Fwd | GTAAAACGACGGCCAGAGTGGCATGGAACAGATT | PANC480_Chr2:125M_td Rev | | CAGGAAACAGCTATGACTGAAAATCAAAAGTATCT |
| PANC480_chr2_164tf_jt2_Fwd | GTAAAACGACGGCCAGTTACCAAAGTTCCCCAGGTG | PANC480_chr2_164tf_jt2_Rev | | CACTTGATTGGGATGAATCG |
| PANC480_chr2_210tf_jt1_Fwd | GTAAAACGACGGCCAGGAGGCAGGCATGGAAAGTTA | PANC480_chr2_210tf_jt1_Rev | | CCCAGAAGGAATGAAGTCCA |
| PANC480_chr2_221tf18_jt1_Fwd | GTAAAACGACGGCCAGAGCAGGCTTTATGCCACATC | PANC480_chr2_221tf18_jt1_Rev | | GGGAAAAGTCTCCCTGGTTC |
| PANC480_chr2_221tf17_jt1_Fwd | GTAAAACGACGGCCAGGCCACATCTTTCCCATTCAA | PANC480_chr2_221tf17_jt1_Rev | | ATCTGACACAAAGGCCCAAG |
| PANC480_Chr2:209M_t Fwd | GTAAAACGACGGCCAGTTAAAGCTTTTGGACTTT | PANC480_Chr2:209M_t Rev | | CAGGAAACAGCTATGACCTGTACTCTGAAAGGATG |
| PANC480_chr2_214t_st1_fwd | GTAAAACGACGGCCAGATTCTACCTGTTCAGGGCCC | PANC480_chr2_214t_st1_rev | | TGTTCAGAGAAGTCTTTGCTCA |
| PANC480_chr2:221M_t Fwd | GTAAAACGACGGCCAGTTCAACTAGGTAGGTCTC | PANC480_chr2:221M_t Rev | | CAGGAAACAGCTATGACTAGCTGGATCTAGGGATT |
| PANC480_chr4_106tf_jt2_Fwd | GTAAAACGACGGCCAGTGAAAGATGCAATGCTCCTG | PANC480_chr4_106tf_jt2_Rev | | CCTCCTCCTGAATTCCTCCT |
| PANC480_Chr4:57M_t_FWD | GTAAAACGACGGCCAGCTGAGCTTATTCTCAGAC | PANC480_Chr4:57M_t_REV | | CAGGAAACAGCTATGACTTCCAACTTCTTTACATC |
| PANC480_chr4:106M_t Fwd | GTAAAACGACGGCCAGCGATCTCAAATCAAACTC | PANC480_chr4:106M_t Rev | | CAGGAAACAGCTATGACGCTACACATATTTCATAA |
| PANC480_chr5_81t_st1_fwd | GTAAAACGACGGCCAGGGGCATACAGGGACAATTCAC | PANC480_chr5_81t_st1_rev | | CCCACCAACCAGAGAGAACT |
| PANC480_chr5_43tf_jt1_Fwd | GTAAAACGACGGCCAGGGTTCCACAGTAACCCAGCA | PANC480_chr5_43tf_jt1_Rev | | CTGTGTGGCTGCTTTCACTG |
| PANC480_chr5_81t_st2_fwd | GTAAAACGACGGCCAGGGGCATACAGGGACAATTCAC | PANC480_chr5_81t_st2_rev | | TGTAAGATGGAGCAGGGACC |
| PANC480_Chr6:28M_d Fwd | GTAAAACGACGGCCAGTTTTCTGCTGATAATTTC | PANC480_Chr6:28M_d Rev | | CAGGAAACAGCTATGACCCTGGATGACATATTTGT |
| PANC480_chr6:25M_td Fwd | GTAAAACGACGGCCAGAGAAAGAAAAGGTAGGAA | PANC480_chr6:25M_td Rev | | CAGGAAACAGCTATGACCTGAATTTACAAATTCGT |
| PANC480_chr6_25id_jt2_Fwd | GTAAAACGACGGCCAGCCACTCCTGGCTTCAAGAAC | PANC480_chr6_25id_jt2_Rev | | GTATGAGGGCCAATTTGTGG |
| PANC480_chr6_27id_fwd1 | GTAAAACGACGGCCAGAGGGACATGTCATAAGCCTCT | PANC480_chr6_27id_rev2 | | TGCGCGTGTTTTAAGAGAGG |
| PANC480_chr8_127tf_fwd1 | GTAAAACGACGGCCAGTAGCTTGATGGGGATGGCAT | PANC480_chr8_127tf_rev1 | | TAACAGGAGAATTGGGCGGT |
| PANC480_Chr9:14M_d Fwd | GTAAAACGACGGCCAGAAAGAAGGAAGGAACCAC | PANC480_Chr9:14M_d Rev | | CAGGAAACAGCTATGACCCAACAAGAGTAAAGGTT |
| PANC480_chr9_78d_st1_fwd | GTAAAACGACGGCCAGGGAACCTCACAAAGTAACTCTGG | PANC480_chr9_78d_st1_rev | | AGGCTCCTTTTGAACACCTTC |
| PANC480_chr9_84t_st2_fwd | GTAAAACGACGGCCAGACACATTCGAAGGAGGCTCA | PANC480_chr9_84t_st1_rev | | AATGAACCACCCTGTCCCAT |
| PANC480_chr18_75i_fwd1 | GTAAAACGACGGCCAGCCACTAGCCTGGCATATCTGA | PANC480_chr18_75i_rev2 | | GGCCCAGATGTCTCACTACA |
| PANC480_chr18_76i_fwd1 | GTAAAACGACGGCCAGTTCATCTATGTCTTTGGTGGCT | PANC480_chr18_76i_rev2 | | CTCCCATCCGAAGAGACAGC |
| PANC504_chr3_60d_jt1_Fwd | GTAAAACGACGGCCAGACACCCCCACCAACTGTAGA | | PANC504_chr3_60d_jt1_Rev | GGCTATACATACCTGCACAGCA |
| PANC504_chr4_21d_st1_fwd | GTAAAACGACGGCCAGAGGATATGTGGAAAGCGCTCT | | PANC504_chr4_21d_st1_rev | TGCATGGCTTCTTCTACAAGTG |
| PANC504_chr4_21td_fwd2 | GTAAAACGACGGCCAGCACATCACATTTGCAGGGGA | | PANC504_chr4_21td_rev1 | GGAACATTGCTCCCCATTCC |
| PANC504_chr4_66td_fwd1 | GTAAAACGACGGCCAGCGTTTCCCAACTAAATGCAGA | | PANC504_chr4_66td_rev1 | TCTTGGGATCATCCTTGACA |
| PANC504_chr4_59i_fwd1 | GTAAAACGACGGCCAGTGGCCCTTATCCCTTCTTTT | | PANC504_chr4_59i_rev1 | CGACCTCCTTCCAATCCAGT |
| PANC504_chr4_2t_fwd1 | GTAAAACGACGGCCAGGGGGACTTGGCTATTTCACA | | PANC504_chr4_2t_rev1 | CTCGTCAGAACCAACGGTCT |
| PANC504_chr4_59t_st1_fwd | GTAAAACGACGGCCAGACTTCCCAGTCAGTGTGTACA | | PANC504_chr4_59t_st1_rev | GCAGGCAAACAGGAACAGAA |
| PANC504_chr6_26d_jt2_Fwd | GTAAAACGACGGCCAGAAGCCCAGGAATTCAAGACC | | PANC504_chr6_26d_jt1_Rev | GTGACAGCGAGTCAGACGTT |
| PANC504_chr7_68d_fwd1 | GTAAAACGACGGCCAGTGGTACAGTTGGTTGATAACACA | | PANC504_chr7_68d_rev1 | AAGTGGAAGAGGTGAAGGGT |
| PANC504_chr7_96d_fwd1 | GTAAAACGACGGCCAGGAGTCCGGGCATTGTACAAG | | PANC504_chr7_96d_rev1 | GGTTTTGTGGCTTCTTGCAT |
| PANC504_chr8_64d_st1_fwd | GTAAAACGACGGCCAGTGCATTTGACGCGCTTGATA | | PANC504_chr8_64d_st1_rev | AAGACGATCGAGACCATCCC |
| PANC504_chr8_145tf_fwd1 | GTAAAACGACGGCCAGCCCCTGATCAGCGTCAAATT | | PANC504_chr8_145tf_rev1 | GCTTTGTTTTCCAGTGCCTG |
| PANC504_chr9_20t_st1_fwd | GTAAAACGACGGCCAGGGGAGGACGCTTCAGAGAAA | | PANC504_chr9_20t_st1_rev | TCTTGAGGAAGGGAGAAACACA |
| PANC504_Chr9:24M_t_Fwd | GTAAAACGACGGCCAGACTTTAGTAATATGTTT | | PANC504_Chr9:24M_t_Rev | CAGGAAACAGCTATGACCTAAGGCAAACAACACTG |
| PANC504_chr11_42t_fwd1 | GTAAAACGACGGCCAGGTCTGTGCTGTCCCTCCTGT | | PANC504_chr11_42t_rev1 | TCCATGGGCACTAGAAGAGC |
| PANC504_chr12_96td_jt1_Fwd | GTAAAACGACGGCCAGAACCCCAACGATCAATTCAC | | PANC504_chr12_96td_jt1_Rev | GCCCTGAGCAATCCTATCTG |
| PANC504_chr12_88t_st1_fwd | GTAAAACGACGGCCAGCACAAAGCCCACACCATGAA | | PANC504_chr12_88t_st1_rev | ACGGGTTGAATGGATTGGTG |
| PANC504_chr14_59t_st1_fwd | GTAAAACGACGGCCAGGGCTCATTCGACTCACTTCC | | PANC504_chr14_59t_st1_rev | GGAGGAATCAGTCTACCCAATT |
| PANC504_chr16_73t_st1_fwd | GTAAAACGACGGCCAGGCCACACATTGTCTCATCCA | | PANC504_chr16_73t_st1_rev | CCAGAAAGGTGAATGCTGTCA |
| PANC504_chr16_75t_fwd2 | GTAAAACGACGGCCAGGGGTTCAAGCAGTTCTCCTG | | PANC504_chr16_75t_rev2 | TCAAACTTCAGCTGGGAACC |
| PANC504_chr17_63t_st1_fwd | GTAAAACGACGGCCAGAATGCAGTGGGGTGAACAAC | | PANC504_chr17_63t_st1_rev | CATGGAGAAACAGGCGAGTG |
| PANC504_chr17_64t_st1_fwd | GTAAAACGACGGCCAGCACCCATTTCTAGTGCTGCC | | PANC504_chr17_64t_st1_rev | CTGGAGAGGCATGGAGAGTT |
| PANC504_Chr17:39M_d Fwd | GTAAAACGACGGCCAGAGTAGGGGTAGAGGACAG | | PANC504_Chr17:39M_d Rev | CAGGAAACAGCTATGACTGTGTGGTTCAGTATATC |
| PANC504_chr17_50id_fwd1 | GTAAAACGACGGCCAGGGAAGTGCAGGCAAAATGAT | | PANC504_chr17_50id_rev1 | TAGCAAGCACCACCTCCTCT |
| PANC504_chr17_66i_fwd1 | GTAAAACGACGGCCAGTGGTCTTCTTTCAAGGTTTGCC | | PANC504_chr17_66i_rev1 | ATAGGTGGTCATTCGAGGGC |
| PANC504_Chr18:50M-1_n1 Fwd | GTAAAACGACGGCCAGAAGCTCTTGAAGACATAA | | PANC504_Chr18:50M-1_n1 Rev | CAGGAAACAGCTATGACATTCCAAAGCCATGCTAA |
| PANC504_Chr18:50M Fwd | GTAAAACGACGGCCAGAGTCAAAGGCCCTCCTCT | | PANC504_Chr18:50M Rev | CAGGAAACAGCTATGACTCCAGCCTCAGACAGAAC |
| PANC504_Chr18:48M Fwd | GTAAAACGACGGCCAGTACCATAGGATGCTTAAC | | PANC504_Chr18:48M Rev | CAGGAAACAGCTATGACTTCAGCCCAGATCCCTAA |
| PANC504_Chr22:30M Fwd | GTAAAACGACGGCCAGGTCCCAGCTACTTGGGAG | | PANC504_Chr22:30M Rev | CAGGAAACAGCTATGACAAGTCAGATCACCTTCAT |
| PANC1002Chr1:74M_d Fwd | GTAAAACGACGGCCAGGGAAACTTCATAAACATT | | PANC1002Chr1:74M_d Rev | CAGGAAACAGCTATGACGTATTTCTCCAACCTATA |
| PANC1002_chr1_72d_jt2_Fwd | GTAAAACGACGGCCAGTTAGGGAGGCAAATCAACCA | | PANC1002_chr1_72d_jt2_Rev | TTTGCTGCAGCTAGCCATTT |
| PANC1002_chr1_72id_fwd2 | GTAAAACGACGGCCAGAATTGTGCCCTGACCATGC | | PANC1002_chr1_72id_rev2 | GAGAGACAGAGACAGAGGTGA |
| PANC1002Chr2:5M_d Fwd | GTAAAACGACGGCCAGGGCGTTCCTTGGGGTTCA | | PANC1002Chr2:5M_d Rev | CAGGAAACAGCTATGACTCATCCAAATCTACTTTC |
| PANC1002Chr2:74M_d Fwd | GTAAAACGACGGCCAGGAAATGATGTCTGGAGGA | | PANC1002Chr2:74M_d Rev | CAGGAAACAGCTATGACTGAGGAAGTGAAAACATT |
| PANC1002Chr2:156M_d Fwd | GTAAAACGACGGCCAGTTCTCTGTTGAGGTTGAC | | PANC1002Chr2:156M_d Rev | CAGGAAACAGCTATGACGCTCTTTTCTTTTTCTTT |
| PANC1002_Chr3:69M Fwd | GTAAAACGACGGCCAGGTCAATATTGAAAGAAGG | | PANC1002_Chr3:69M Rev | CAGGAAACAGCTATGACACCCAGTTAACATCACAA |
| PANC1002_Chr4:178M Fwd | GTAAAACGACGGCCAGTATAGCCATCATAGCATA | | PANC1002_Chr4:178M Rev | CAGGAAACAGCTATGACGCACCTACCTCACCTGCA |
| PANC1002Chr5:27439M_d Fwd | GTAAAACGACGGCCAGAAGCTGCAGATCTTCACG | | PANC1002Chr5:27439M_d Rev | CAGGAAACAGCTATGACTTCTGTAATTCTACAAGA |
| PANC1002Chr5:27824M_d Fwd | GTAAAACGACGGCCAGGTAATATATTTAAAGATT | | PANC1002Chr5:27824M_d Rev | CAGGAAACAGCTATGACAAGATGGTGAAGAATTAG |
| PANC1002Chr5:115M_Hd Fwd | GTAAAACGACGGCCAGCTCTAGATCTGGATGAGG | | PANC1002Chr5:115M_Hd Rev | CAGGAAACAGCTATGACGAAGCAGGGTTTTCTGCA |
| PANC1002Chr5:26M_d Fwd | GTAAAACGACGGCCAGAATATGGAAGATACTAAT | | PANC1002Chr5:26M_d Rev | CAGGAAACAGCTATGACGTAAATGTCATATTGTGA |
| PANC1002_chr5_22t_st1_fwd | GTAAAACGACGGCCAGCCAAATATGAAAGCCCCAAA | | PANC1002_chr5_22t_st1_rev | GGGGTTCAGAACTTCAGTGG |
| PANC1002_Chr6:81M Fwd | GTAAAACGACGGCCAGTCTTCTGTGTCGCTCACG | | PANC1002_Chr6:81M_n1 Rev | CAGGAAACAGCTATGACTATGATCACCTTGTATAA |
| PANC1002_chr7_3d_jt1_Fwd | GTAAAACGACGGCCAGGTGAATTTCCTGGGGTTCAG | | PANC1002_chr7_3d_jt1_Rev | ATGGATTGGGTGTCCAGAAA |
| PANC1002Chr7:344M_d Fwd | GTAAAACGACGGCCAGTGATGGCACAAAGGAAAA | | PANC1002Chr7:34M_d Rev | CAGGAAACAGCTATGACAATGGGAAAGATATATAA |
| PANC1002_chr7_111d_st1_fwd | GTAAAACGACGGCCAGGGGTTGCAGTCTTCCTTGTC | | PANC1002_chr7_111d_st1_rev | TGGGAGAAGACCCAGCTAAA |
| PANC1002_Chr8:123M Fwd | GTAAAACGACGGCCAGTACCAATTACATGTGAGG | | PANC1002_Chr8:123M Rev | CAGGAAACAGCTATGACCCTCCAAATACCATCCCA |
| PANC1002_Chr8:138M_n1 Fwd | GTAAAACGACGGCCAGTGTGATAGGCTAAATAAT | | PANC1002_Chr8:138M_n1 Rev | CAGGAAACAGCTATGACTTCCTGTCCAGCATTCAC |
| PANC1002_chr8_51d_st1_fwd | GTAAAACGACGGCCAGAGATGGAGAAGGGAATGCAA | | PANC1002_chr8_51d_st1_rev | TGCGTTGTTATCATACTGTGC |
| PANC1002_chr9_21t_fwd1 | GTAAAACGACGGCCAGATTAGCCCCTGGAAAGCAGT | | PANC1002_chr9_21t_rev2 | ACATGCCGTACAAGTCATCC |
| PANC1002_chr9_21995t_st1_fwd | GTAAAACGACGGCCAGATTGTGCAGAAGCCAGTCCT | | PANC1002_chr9_21995t_st1_rev | GGGATGGGGAAAGAGAAGTC |
| PANC1002_chr12_28i_jt1_Fwd | GTAAAACGACGGCCAGCCCATTGCAAGCCTACAGTT | | PANC1002_chr12_28i_jt1_Rev | TCCCTGAGAAAGTCCTGGTTT |
| PANC1002Chr12:86M_d Fwd | GTAAAACGACGGCCAGATCTTTCTCTTACCCTAC | | PANC1002Chr12:86M_d Rev | CAGGAAACAGCTATGACTGTTAACTAGAATAA |
| PANC1002_chr13_53d_fwd1 | GTAAAACGACGGCCAGGGGACAGTAGAGGCATCAGA | | PANC1002_chr13_53d_rev2 | GACAAAGTGGCATGGCATGA |
| PANC1002Chr13:82M_d Fwd | GTAAAACGACGGCCAGAAATGTTTTTGAAGTTCA | | PANC1002Chr13:82M_d Rev | CAGGAAACAGCTATGACTTCCCTGCAATGGAGGGC |
| PANC1002Chr13:95M_d Fwd | GTAAAACGACGGCCAGATCATTTTATCTTCAATT | | PANC1002Chr13:95M_d Rev | CAGGAAACAGCTATGACGAAAAGGCAAAACCACAA |
| PANC1002_chr17_11tf_fwd1 | GTAAAACGACGGCCAGGCTTGTGGGAAATGCAGAAT | | PANC1002_chr17_11tf_rev2 | CACCAAGCCATTCATGAGGG |
| PANC1002_chr17_12t_st1_fwd | GTAAAACGACGGCCAGCTTCCCCTCCCTAGTTGACC | | PANC1002_chr17_12t_st1_rev | GAAGGGGGAAAAGGGTGATA |
| PANC1002_Chr18:48M Fwd | GTAAAACGACGGCCAGGCATTGTAGATTCATACA | | PANC1002_Chr18:48M_n1 Rev | CAGGAAACAGCTATGACATTGGCTGGTGGGCACAC |

**^*^**Primers were named by their target cell line (e.g. “Panc480”), chromosome location (e.g. “chr1”) followed by either the first few numbers of the coordinates in the thousands (e.g. “550”) or the millions (e.g. “53M”).

**^#^**M13F sequence was adapted to forward primers for Sanger sequencing.

**Table 5. Primers for Cas9-mApple, Cas9-mNeonGreen, and dCas9-EGFP plasmid constructions and validations.**

| **Name** | **Sequence** | **Purpose** |
| --- | --- | --- |
| Vector forward | ctccaccggcggcatggacgagctgtacaagcatcatcac | Cas9-mApple Gibson assembly |
| Vector reverse | ccatgttattctcctcgcccttgctcaccatggtggcgac |  |
| Insert forward | gggcgaggagaataacatggccatcatcaaggagttcatg |  |
| Insert Reverse | cgtccatgccgccggtggagtggcggccctcggcgcgttc |  |
| mCherry-F | CCCCGTAATGCAGAAGAAGA | Cas9-mApple insertion validation |
| WPRE-R | CATAGCGTAAAAGGAGCAACA |  |
| mNG_fwd-2 (insert) | gtgcccgtcagtgggcag | Cas9-mNeonGreen Gibson assembly |
| mNG_rev-2 (insert) | agcagagagaagtttgttgcgccggatc |  |
| EGFP_vec_fwd (vector) | gcaacaaacttctctctgct |  |
| EGFP_vec_rev (vector) | gggcgatgtgcgctctgccc |  |
| Vector forward | gtacgagacacggatcgacctgtctcagctgggaggcgacaagcgacctgccgccacaaa | dCas9-EGFP Gibson assembly |
| Vector reverse | ctgtgttctggcggcaaacccgttgcgaaaaagaacgttcacggcgactactgcacttat |  |
| Insert forward | gaacgttctttttcgcaacgggtttgccgccagaacacaggaccggtgccgcccaccatg |  |
| Insert Reverse | gtcgcctcccagctgagacaggtcgatccgtgtctcgtacaggccggtgatgctctggtg |  |
| D10 Forward | tggctccgcctttttcccga | dCas9-EGFP plasmid validation (primers amplify across both nuclease domains) |
| D10 Reverse | ctcggctgtttctccgctgt |  |
| H840 Forward | gagctgggcagccagatcct |  |
| H840 Reverse | cttggcattcagcagctggc |  |

**Table 6. Primers for Cas9 activity assay.**

| **Name** | **Sequence** | **Purpose** |
| --- | --- | --- |
| i_HPRTc.465_Fwd-2 | AATGATACGGCGACCACCGAGATCTACACTCTTTCCCTACACGACGCTCTTCCGATCTATCATGCTTAGAGGGCCAGATGATATAGATTCC | NGS primers for human cell lines |
| ib_HPRTc.465_Rev-2 | CAAGCAGAAGACGGCATACGAGATATAGCGTCGTGACTGGAGTTCAGACGTGTGCTCTTCCGATCTGGCAAGGAAGTGACTGTAATTATG |  |
| mchrX_52M_Fwd | AATGATACGGCGACCACCGAGATCTACACTCTTTCCCTACACGACGCTCTTCCGATCTTAAGTAGAGTGCTCCACTTTGAAACAGCTG | NGS primers for mouse cell lines |
| mchrX_52M_Rev | CAAGCAGAAGACGGCATACGAGATTCGCCTTGGTGACTGGAGTTCAGACGTGTGCTCTTCCGATCTACACATGCCTCTCCTCTCTCT |  |

**Table 7. Primers involved in multiplex sgRNA vector construction.**

| **Primer name** | **Sequence** | **Purpose** |
| --- | --- | --- |
| Multi_lenti_frag_fwd1 | cccacctcccaaccccgaggggacccagagagggcctatttc | amplification of sgRNA cassettes |
| Multi_lenti_rev_2 | gggaaataggccctctctgggtcgaaaaaagcaccgactcggtgccactt |  |
| multiplex-BsrGI-fwd | tatcgttgTGTACAaggcagggatattcaccatt | amplification of LOH array out of lentiGuide |
| multiplex-MreI-rev | tatcgttgCGCCGGCGaattgtggatgaatactgcc |  |
| lentiC_vecfwd-MreI | tatcgttgCGCCGGCGgaattcgctagctaggtcttg | linearization of lentiCRISPRv2-puro |
| lentiC_vecrev-BsrGI | tatcgttgTGTACAccaaactggatctctgc |  |
| lentiG_vecfwd-MreI | tatcgttgCGCCGGCGgagacaaatggcagtattcatc | linearization of lentiGuide-puro |
| lentiG_vecrev-BsrGI | tatcgttgTGTACActctattcactatagaaagtacagcaaaaactattcttaaacc |  |
| Stitch_fragFwd | agggatattcaccattatcgtcgtttcagacccacct | Gibson Assembly of LOH-7 partial assemblies |
| Stitch_fragRev | gggttgggaggtgggtctgactcaagatctagttacgccaagct |  |
| Stitch_vectorFwd | tggcgtaactagatcttgagtcagacccacctcccaaccc |  |
| Stitch_vectorRev | gggaggtgggtctgaaacgacgataatggtgaa |  |
| Mulitplex_lenti_fwd1 | aggcagggatattcaccatt | Construct validation |
| Mulitplex_lenti_rev2 | aattgtggatgaatactgcc |  |
| 480LOHG1_fwd | GGAATCATCTTCACAGTTGT |  |
| 480LOHG1_Rev | ACAACTGTGAAGATGATTCC |  |
| 480LOHG4_fwd | CTAATGTATGACTGAAAGCT |  |
| 480LOHG4_Rev | AGCTTTCAGTCATACATTAG |  |
| 480LOHG5_fwd | GAGGTGTCTAAACCATGACA |  |
| 480LOHG5_Rev | TGTCATGGTTTAGACACCTC |  |
| pFH6-seq_fwd | ctgcaggtcgaccatatggg |  |
