## Supplementary material for "CRISPR-Cas9 for selective targeting of somatic mutations in pancreatic cancers": figure S: Supplementary figure S1-6.pdf

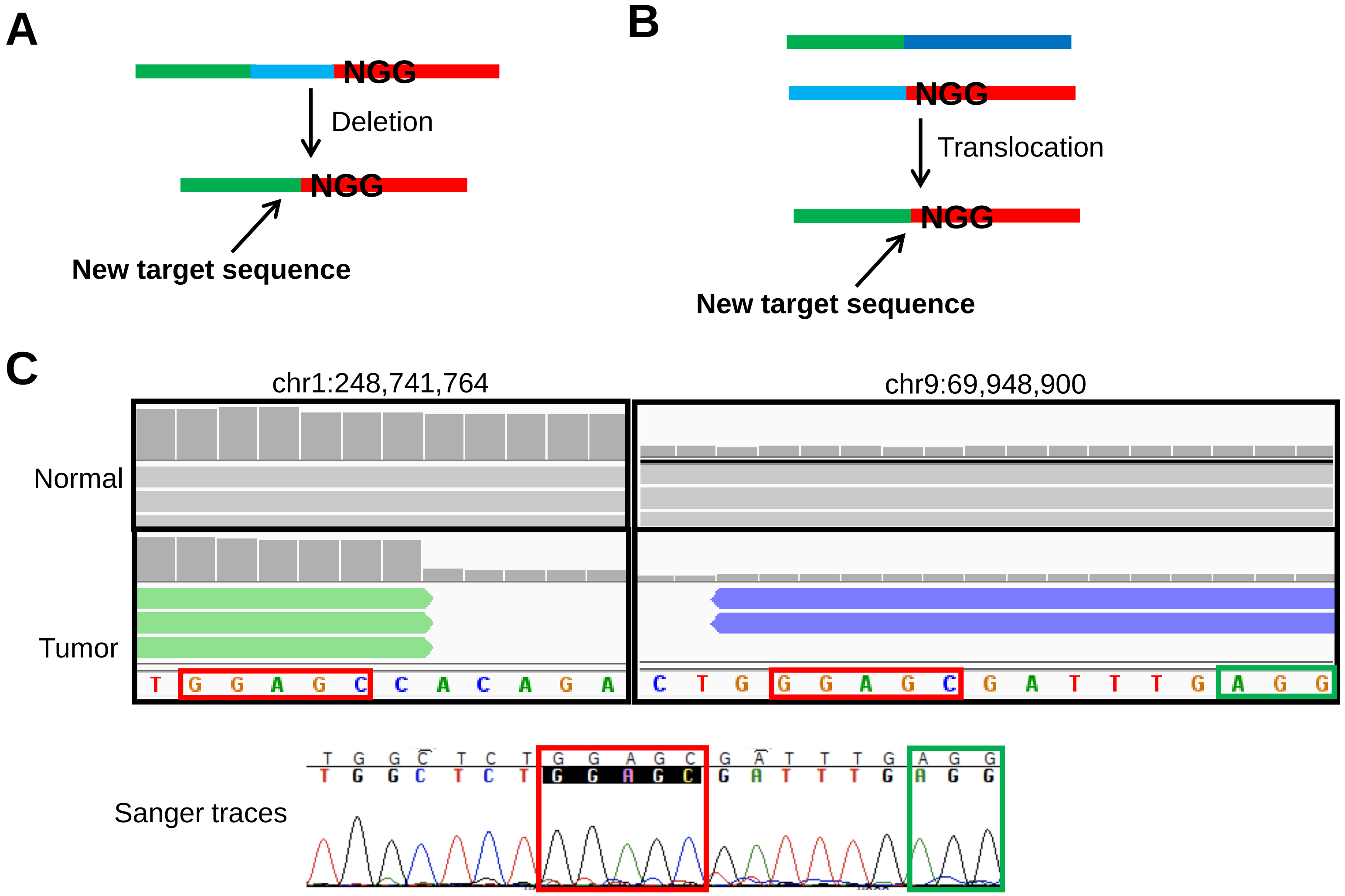


**Figure S1. Structural variants create novel CRISPR-Cas9 target sites.** Structural variants, such as (A) deletion and (B) translocation, could give rise to novel target sequence if the new junction is in the proximity of an existing NGG PAM (shown) or creates a new PAM (not shown). For example, (C) a chr1:chr9 translocation in Panc480 gave rise to a novel breakpoint that was in proximity of an existing AGG PAM (green box). This breakpoint was characterized by a 5bp GGAGC microhomology at its junction (red box).

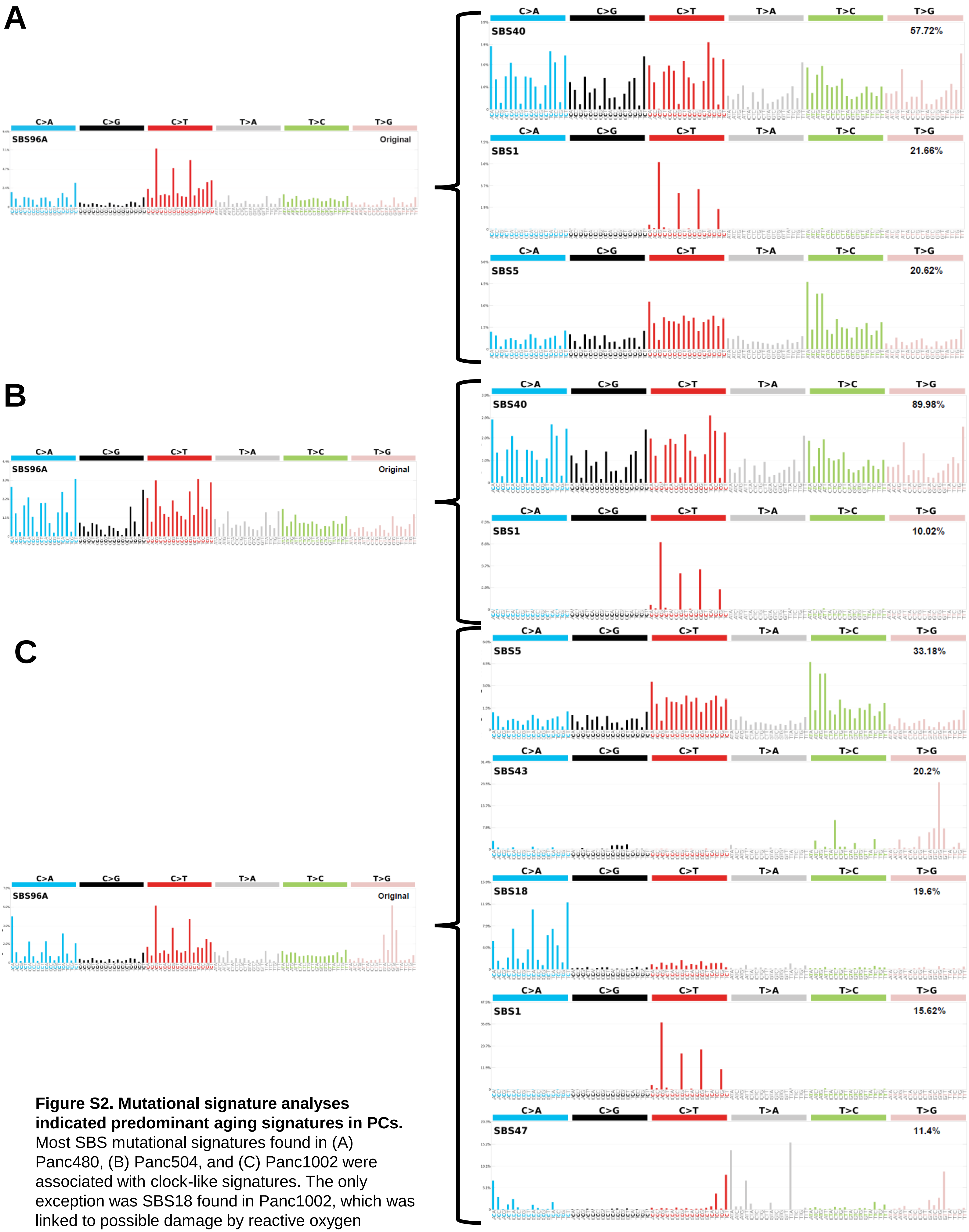


**Figure S2. Mutational signature analyses indicated predominant aging signatures in PCs.** Most SBS mutational signatures found in (A) Panc480, (B) Panc504, and (C) Panc1002 were associated with clock-like signatures. The only exception was SBS18 found in Panc1002, which was linked to possible damage by reactive oxygen species. Y-axis is the percentage of SBS.

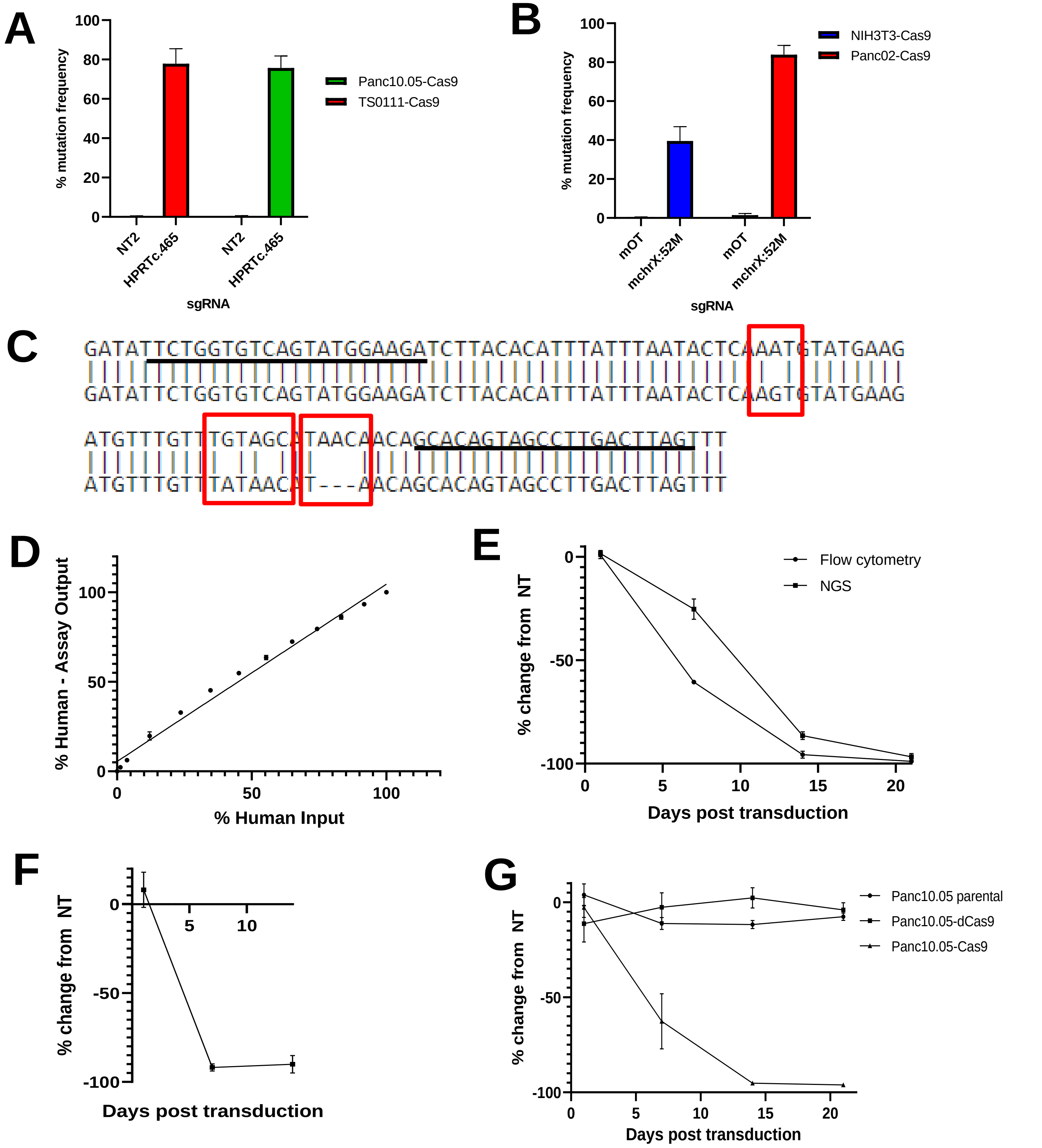


**Figure S3. Human cell line-specific toxicity was reproducible across different combinations of mouse-human co-cultures, and this selective cell elimination required the presence of both Cas9 and human-specific sgRNA.** (A-B) Cas9 activity assay was performed on (A) PC Cas9-expressing cell lines (Panc10.05, TS0111) and (B) mouse Cas9-expressing cell lines (NIH 3T3 and Panc02) to quantify mutation frequency at the *HPRT1* gene locus. (C) Alignment of the mouse and human *RC3H2* orthologs shows differences of a 3bp indel and 3 SNPs between the two species, highlighted by red boxes. PCR primer sequences are underlined. (D) Sensitivity and accuracy of the mouse-human NGS assay was validated by deep sequencing known mixes of mouse and human DNA. Pearson r = 0.9941, *P<0.0001*, N=3. (E) TS0111 and NIH 3T3 Cas9-expressing cell lines were co-cultured and transduced with 230F(12). Shown are the changes in TS0111 cell population over time by flow cytometry (based on fluorophore expression) and the mouse-human NGS assay. (F) Panc10.05 and Panc02 Cas9-expressing cell lines were also co-cultured and transduced with the same sgRNA, in which the change in Panc10.05 cell population was measured by flow cytometry. Panc02 is a KPC-derived mouse cell line. (G) NIH 3T3 Cas9-expressing cell line was co-cultured with either Panc10.05 parental, dCas9-expressing cell line, or Cas9-expressing cell line, and transduced with 230F(12), in which the change in NIH 3T3 cell population was measured by flow cytometry. For E-G, error bars indicate mean ± SEM; N=3.

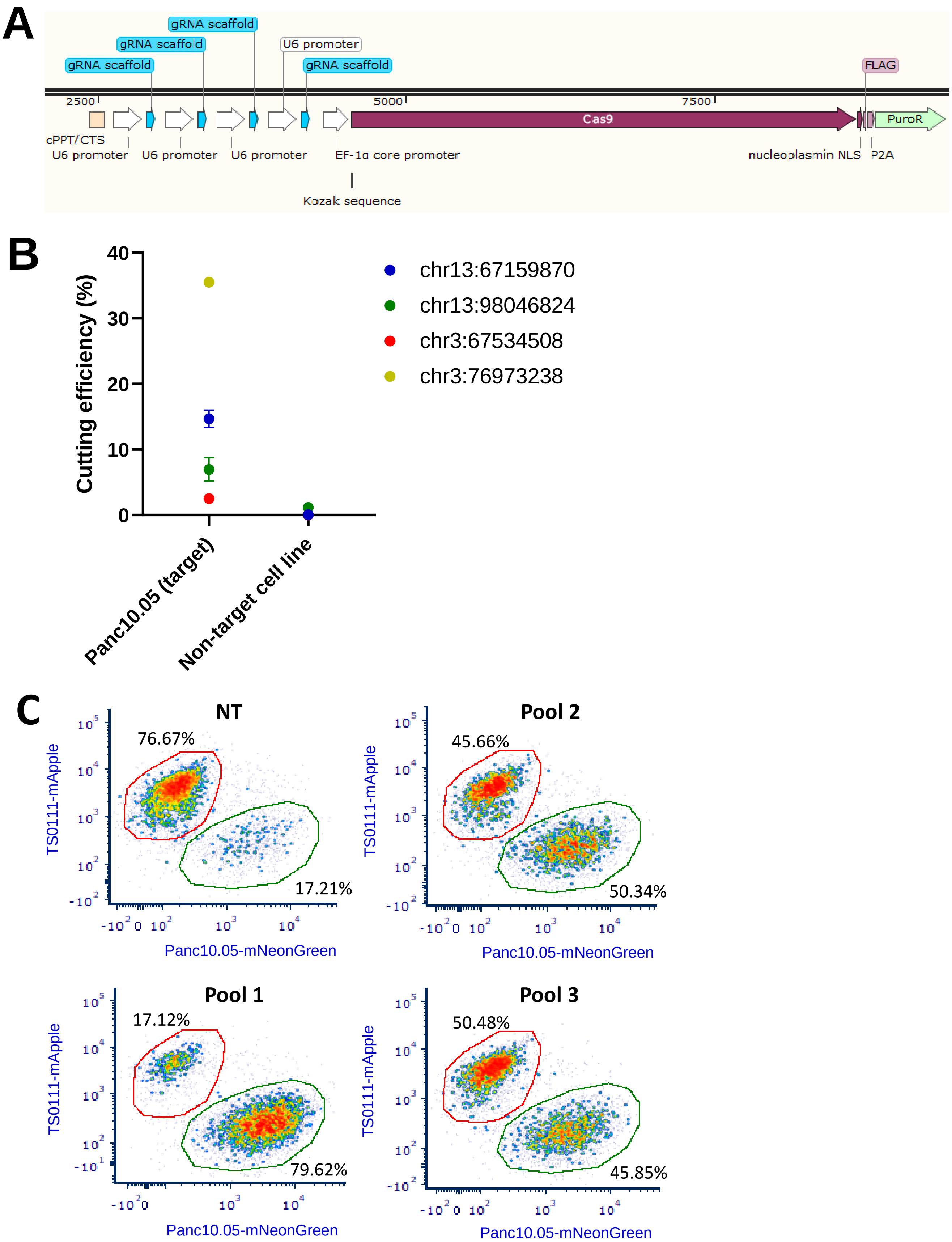


**Figure S4. Panc10.05 quad produced on-target mutations, and TS0111-specific sgRNAs produced selective toxicity.** (A) Illustration of Panc10.05 quad vector. The vector was designed to express 4 sgRNAs targeting Panc10.05 simultaneously. Diagram was generated by SnapGene. (B) Panc10.05 quad was transduced into Panc10.05 and a negative control cell line (Panc480), and mutation frequency was quantified 21 days post transduction using deep sequencing and CRISPResso2. N=1/2; mean ± SEM are shown. One of the loci in Panc10.05 quad has a noncanonical germline PAM and cut Panc480 at a 1.16% mutation frequency. (C) Co-cultures of TS0111 (labeled with mApple) and Panc10.05 (labeled with mNeonGreen) were treated with three different pools of TS0111-specific sgRNAs (9 sgRNA per pool) along with equivalent doses of non-targeting sgRNA controls (NT; MOI90), and flow cytometry was performed to quantify cells that were positive for either mNeon-Green or mApple. Flow cytometry analyses of one replicate on day 14 post transductions were shown. NT and Pool 1 data were also displayed in Figure 3E.

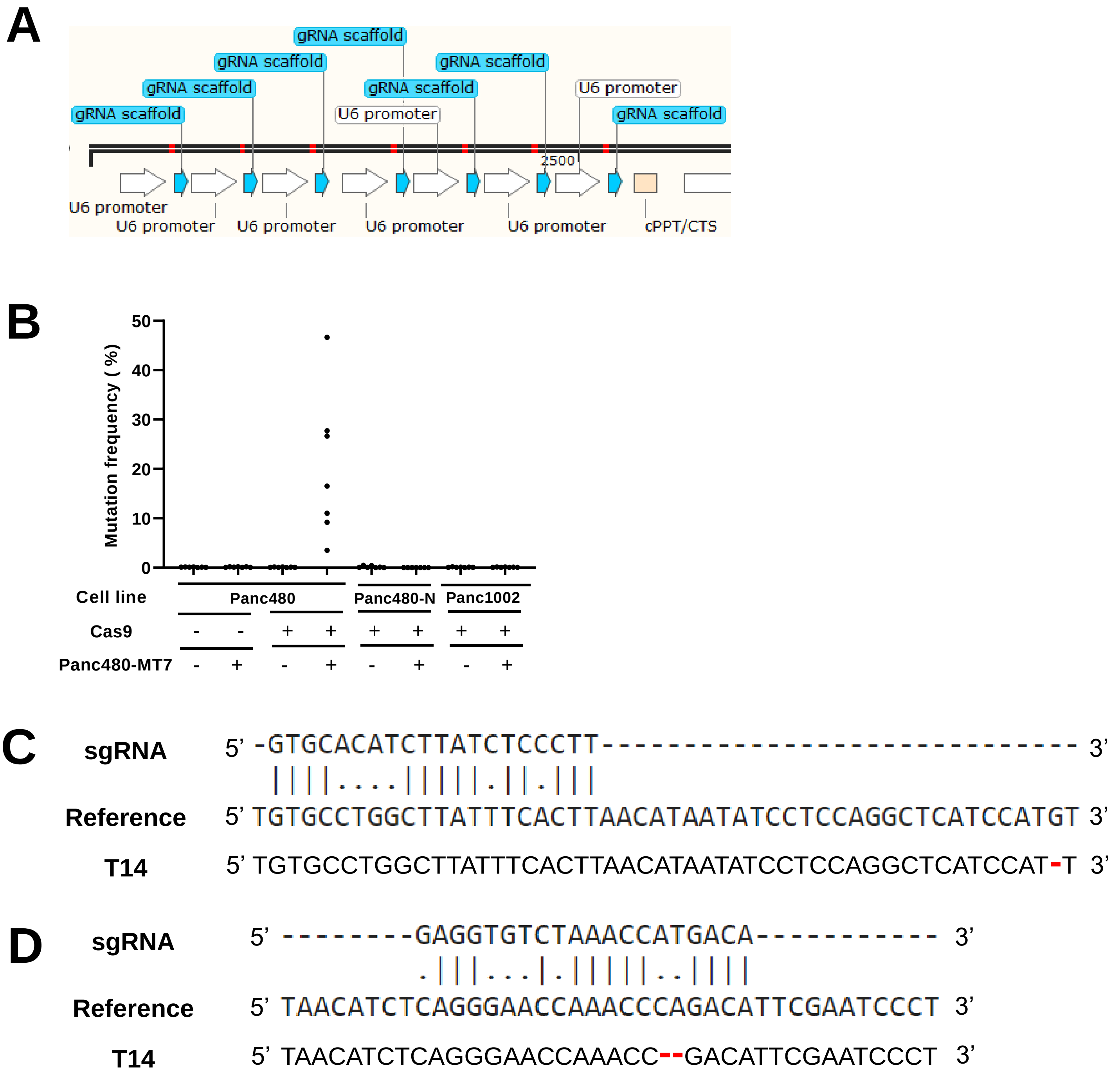


**Figure S5. Absence of off-target activity of Panc480-specific sgRNAs.** (A) Illustration of Panc480-MT7 vector. The vector was designed to express 7 sgRNAs (red line indicates 20bp spacer) targeting Panc480 simultaneously. Diagram was generated by SnapGene. (B) Mutation frequency at 7 Panc480-specific target sites in Panc480 parental, Panc480 Cas9-expressing, Panc480 patient's Cas9-expressing lymphoblasts (normal cell line, Onc3286, indicated as Panc480-N), and Panc1002 Cas9-expressing (negative control) cell lines after treatment with NT (-) or Panc480-MT7 (+) multiplex sgRNA vector. (C) Pairwise sequence alignments (EMBOSS Needle) among Panc480-MT7 chr8:29032916 sgRNA sequence (DNA bases are shown), the region surrounding chr8:7530860 in the hg19 human reference genome that has the lowest number of potential mismatches (6bp), and the actual sequence in Panc1002 T14. Red dash indicates deletion. (D) Pairwise sequence alignments among chrX:3982448 sgRNA sequence (DNA bases are shown), the region surrounding chr15:24671815 in hg19 that has the lowest number of potential mismatches (7bp), and the actual sequence in Panc1002 T14. Red dash indicates deletion.

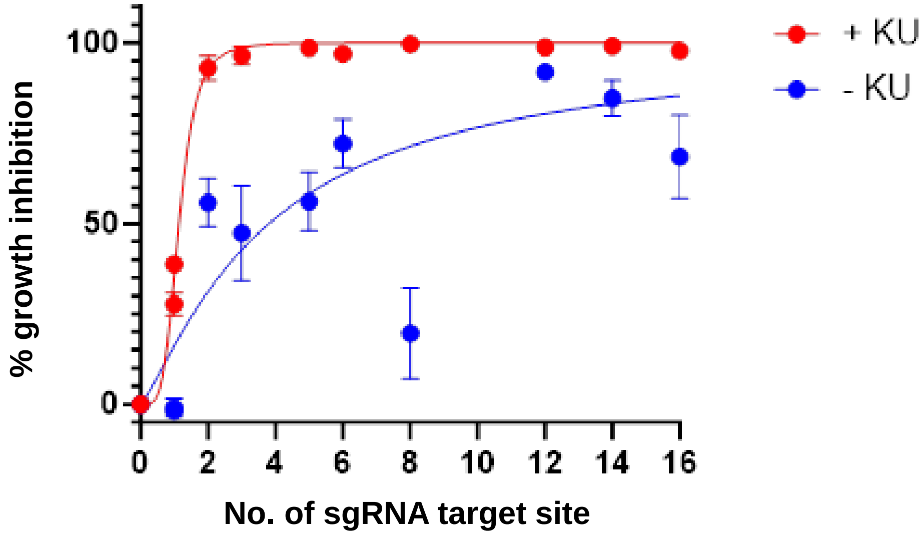


**Figure S6. DSB repair inhibitor increased growth inhibition of multitarget sgRNAs.** sgRNAs with 0-16 target sites in the human genome were transduced into TS0111 and treated with either DMSO (-KU) or 1uM DNA-PK/PI3-K inhibitor, KU-0060648 (+KU). Growth inhibition was assessed using alamarBlue cell viability assay.
