## Supplementary material for "CRISPR-Cas9 for selective targeting of somatic mutations in pancreatic cancers": figure S: Supplementary Materials and Methods.docx

*SV target validation and sgRNA design*

Genomic DNA from tumors and corresponding normal tissues of Panc480, Panc504, and Panc1002 were used for high-density SNP microarray and whole genome sequencing (WGS) as previously described (1,2). A list of structural variants (SVs) were compiled from SVs previously published in Norris *et al.* (2015) (2). Additional SVs were discovered by using Trellis (3), an SV caller on WGS data via tumor-normal (T-N) subtraction. SVs that were present in normal based on IGV (4) visual inspection were further eliminated from the list. Primers were designed to PCR amplify across breakpoints and sent for Sanger sequencing (Primers Table 4). Among the validated ones, we designed potential sgRNA sequences in which either the PAM spanned across the breakpoint junction or at least 4bp of the sgRNA sequence crossed the junction. Then, we entered the sequences into CRISPOR and selected candidates that have >50 specificity score (5).

*WES target identification and sgRNA design*

Genomic DNA from tumors and corresponding normal tissues of Panc480, Panc504, and Panc1002 were whole exome sequenced and variants called as previously described (1). Mutations were inspected to include novel Cs that were adjacent to an existing C or novel Gs that were adjacent to an existing G after T-N subtraction. The resulting list of mutations was put through CRISPOR and the ones that produced sgRNAs with >50 specificity score in CRISPOR were subsequently examined for their variant allele frequencies (VAFs) (5).

*Cas9-mApple plasmid construction*

pLentiCas9-T2A-GFP was a gift from Roderic Guigo & Rory Johnson (Addgene plasmid # 78548, (6)) and mApple-N1 was a gift from Michael Davidson (Addgene plasmid # 54567, (7)). Primers were designed to amplify the vector from pLentiCas9-T2A-GFP and mApple insert from mApple-N1 using Q5 Hot Start High-Fidelity polymerase (NEB) according to the manufacturer’s protocol (Primers Table 5). PCR products were subjected to gel electrophoresis with 0.8% agorose gel at 150V for 2 hours. Gel extraction was performed with QIAquick Gel Extraction Kit (QIAGEN) according to the manufacturer’s protocol to purify the vectors and inserts. Then, Gibson assembly was performed with a 2:1 ratio of insert:vector using Gibson Assembly Master Mix (NEB) and an incubation time of 1 hour at 50°C. The Gibson product was transformed into NEB 5-alpha Competent E. coli according to the manufacturer’s protocol and were selected by both carbenicillin and ampicillin. Plasmids were extracted from ampicillin-resistant clones using QIAprep Spin Miniprep kit (QIAGEN) according to the manufacturer’s protocol. Analytical digestion with restriction enzymes (NEB) was performed to verify the identity of the plasmid. Primers were designed to confirm insertion (Primers Table 5). The plasmid was then transfected into 293T cells with Invitrogen Lipofectamine 3000 reagent and P3000 reagent (ThermoFisher) according to the manufacturer’s protocol, and observed under fluorescence microscope for functional validation. The final plasmid has been validated by whole plasmid sequencing (Plasmidsaurus).

*Cas9-mNeonGreen plasmid construction*

pLenti-spCas9-T2A-mNeonGreen-P2A_puro was a gift from Raphael Gaudin (Addgene plasmid #122183). Primers were designed to amplify the vector from pLenti-Cas9-T2A-EGFP and mNeonGreen insert from pLenti-spCas9-T2A-mNeonGreen-P2A_puro using Invitrogen Platinum Superfi II PCR Mastermix (ThermoFisher) according to the manufacturer’s protocol (Primers Table 5). The resulting fragments were digested with DpnI at 37°C for 1 hour and then concentrated by QIAquick Column (QIAGEN). Gibson assembly was performed at a 1:1 molar ratio, using 0.018 pmol per fragment, with the Invitrogen Gene Art Gibson Assembly Kit (ThermoFisher). 2uL of the Gibson assembly reaction was transformed into 25uL OneShot Stbl3 competent cells (ThermoFisher, USA) by heat shock following the manufacturer’s instructions. Correct constructs were verified by analytical digest and Sanger sequencing (Primers Table 5). The final plasmid has been validated by whole plasmid sequencing (Plasmidsaurus).

*mNeonGreen-bc plasmid construction*

Barcoded plasmids were created through a restriction-ligation insertion of a barcode sequence into a donor plasmid followed by PCR linearization and blunt-end ligation to remove Cas9 from final product. Barcode single-strand oligo inserts were ordered from IDT. Barcode oligos were comprised of a spacer sequence, a ClaI recognition sequence, the M13F sequence, an 8-bp barcode sequence, the XhoI restriction site, and another spacer sequence. Oligo inserts were ordered as pairs of reverse complements to create a double-strand final product. Matched oligo pairs were phosphorylated and annealed using T4 ligation buffer and T4 PNK (NEB). Annealed oligos were then digested with ClaI and XhoI to create sticky ends complementary to the destination plasmid and diluted 1:200 in DEPC treated water. The destination plasmids, pLenti_Cas9_T2A_mApple-blasticidin (see Cas9-mApple plasmid construction) and pLenti_Cas9_T2A-mNeonGreen-Blasticidin (see Cas9-mNeonGreen plasmid construction), were also digested with ClaI and XhoI (NEB) for 15 minutes at 37°C. 1uL insert and 50ng vector were then annealed using the Rapid DNA Ligation Kit (ThermoFisher). The ligation mixture was transformed into OneShot Stbl3 competent cells (ThermoFisher) following the manufacturer’s directions. The resultant plasmids were then PCR linearized using 5’-gtgagcaagggcgaggag and 5’-ccatggtggcaccggtc for mNeonGreen and 5’-catggtgagcaagggcg and 5’-gtggcagcgctctagaac for mApple. The linearized PCR products were digested with DPNI and concentrated by QIAquick column (QIAGEN) prior to blunt end ligation of 20ng vector with the DNA Rapid Ligation Kit (ThermoFisher). The final plasmid has been validated by whole plasmid sequencing (Plasmidsaurus).

*dCas9 plasmid construction*

pLentiCas9-T2A-GFP was a gift from Roderic Guigo & Rory Johnson (Addgene plasmid # 78548, (6)) and pZLCv2-3xFLAG-dCas9-HA-2xNLS was a gift from Stephen Tapscott (Addgene plasmid # 106357, (8)). Primers were designed to amplify the vector from pLentiCas9-T2A-GFP and dCas9 insert from pZLCv2-3xFLAG-dCas9-HA-2xNLS using Q5 Hot Start High-Fidelity polymerase (NEB) according to the manufacturer’s protocol (Primers Table 5). PCR products were subjected to gel electrophoresis with 0.8% agarose gel at 150V for 2 hours. Gel extraction was performed with QIAquick Gel Extraction Kit (QIAGEN) according to the manufacturer’s protocol to purify the vectors and inserts. Then, Gibson assembly was performed with a 3:1 ratio of insert:vector using Gibson Assembly Master Mix (NEB) and an incubation time of 1 hour at 50°C. The Gibson product was transformed into NEB 5-alpha Competent E. coli according to the manufacturer’s protocol and were selected by both carbenicillin and ampicillin. Plasmids were extracted from ampicillin-resistant clones using QIAprep Spin Miniprep kit (QIAGEN) according to the manufacturer’s protocol. Analytical digestion with restriction enzymes (NEB) was performed to verify the identity of the plasmid. Primers were designed to PCR and Sanger sequence regions spanning D10 and H840 of dCas9 to validate the mutations on dCas9 (Primers Table 5). The final plasmid has been validated by whole plasmid sequencing (Plasmidsaurus).

*sgRNA-expressing plasmid construction*

lentiGuide-Puro was a gift from Feng Zhang (Addgene plasmid # 52963, (9)) and lentiCRISPRv2 puro was a gift from Brett Stringer (Addgene plasmid # 98290, (10)). Oligonucleotides of sgRNA sequences were ordered from IDT for cloning into both lentiGuide-Puro and lentiCRISPRv2 puro backbones according to Feng Zhang’s Lab Target Guide Sequence Cloning protocol (9,11). The resulting product was transformed into One Shot Stbl3 chemically competent E. coli (ThermoFisher) according to the manufacturer’s protocol and selected with both carbenicillin and ampicillin. Plasmids were extracted from ampicillin-resistant clones using QIAprep Spin Miniprep kit (QIAGEN) according to the manufacturer’s protocol. Analytical digestion with restriction enzymes (NEB) was performed to verify the identity of the plasmids and Sanger sequencing was performed to validate the insertion of sgRNA sequence.

*Lentivirus titer preparation and quantification*

pCMV-VSV-G was a gift from Dr. Bob Weinberg (Addgene plasmid #8454, (12)), pMDLg/pRRE and pRSV-Rev were gifts from Dr. Didier Trono (Addgene plasmid #12251 & #12253, (13)). 2.5ug pCMV-VSV-G, 5ug pMDLg/pRRE, 5ug pRSV-Rev, and 7.5ug transfer plasmids were used along with 50uL Invitrogen Lipofectamine 3000 reagent and 40uL P3000 reagent (ThermoFisher) for transfection into 293T cells on a 10-cm plate (95-99% confluent at transfection). Cell culture and transfection workflows were the same as the manufacturer’s protocol. Upon harvesting and pooling the lenvirus-containing supernatant, the clarified supernatant was concentrated with Lenti-X Concentrator (Takara Bio) by following the manufacturer’s protocol. Lenti-X qRT-PCR titration kit (Takara Bio) was used to quantify an aliquot of the clarified lentiviral supernatant according to the manufacturer’s protocol.

*Cell culture*

Panc10.05, TS0111, Panc480, Panc1002, NIH 3T3, Panc02, Onc3286, and their derivative cell lines were STR profiled and mycoplasma tested before the start of experiments. All cells, except for Onc3286, were maintained in monolayer cultures at 37˚C and 5% CO_2_. The culture medium consisted of 1X DMEM, 10% fetal bovine serum, 2mM L-glutamine, and 1X antibiotic antimycotic solution (Sigma; contains 100u penicillin, 100ug streptomycin, and 0.25ug amphotericin B). Onc3286 was maintained in a suspension culture at 37˚C and 5% CO_2_. The culture medium consisted of 1X RPMI 1640, 20% heat-inactivated bovine calf serum, 2mM L-glutamine, and 1X antibiotic antimycotic solution (Sigma).

*Fluorescent cell line construction*

Cells were seeded at 50% confluence for 24 hours before the media was replaced to contain 10ug/mL of polybrene. Lentivirus of fluorophore-expressing plasmids were added into the media at MOI 0.01 and transduction took place for 18-20 hours. The media was then removed, washed once with PBS, and replaced with normal media. After 24 hours, the media was replaced with media that contained 5ug/mL blasticidin for a 7-day selection. The cells were then sent to the SKCCC Flow Cytometry Core or SKCCC High Parameter Flow Core for fluorescence activated cell sorting using BD FACSAria II or BD Fusion sorter, respectively, to sort for cells with the optimal fluorescence intensity. The sorted cells were cultured in the presence of blasticidin selection and subjected to STR profiling and mycoplasma testing. Fluorescence microscopy was performed to verify the presence of fluorescent markers before experiments were carried out on these cell lines.

*Next generation sequencing (NGS) of amplicons*

PCR was performed with primers containing partial Illumina adapter sequences to generate amplicons. Either NEBNext High-Fidelity 2X PCR Master Mix (NEB) or Platinum SuperFi II PCR Master Mix (Thermo Fisher) was used for PCR preparations, and thermocycling conditions were set based on manufacturers’ suggestions. Amplicons were purified using QIAGEN MinElute PCR purification kit based on manufacturer’s protocol. Purified PCR products were sent to Azenta for Amplicon-EZ service, in which 2x250bp sequencing was performed to provide ~50,000 reads per sample. FASTQ files were obtained for further analysis.

*Cas9 activity assay*

Cells were transduced with sgRNAs targeting *HPRT1* gene to induce mutations, which could be functionally screened via 6-thioguanine (6-TG) positive selection. For human, the sgRNA used was HPRTc.465 (designed via CRISPOR (5)) and non-targeting control was NT2 (14); for mouse, it was mchrX:52M with mchrX:53M as an off-target control, both designed via CRISPOR (Primers Table 6). Target site was PCR amplified and sent for NGS (Primers Table 6). Mutation frequency of target site was quantified using CRISPResso2 pipeline (15).

*Multiplex sgRNA vector construction*

Individual sgRNA targeting novel PAMs were obtained as ssDNA oligos from IDT and cloned into lentiGuide-puro (Addgene #52963), lentiCRISPRv2-puro (Addgene #98290), or lentiCRISPRv2-hygro (Addgene #98291) lentiviral expression backbones per the protocol previously published by the Zhang Lab (9–11). The U6 promoter, sgRNA spacer sequence, and sgRNA scaffold, referred to here as cassettes, were then PCR amplified off each backbone (Primers Table 7). For multiplexing, the backbone containing the first sgRNA was linearized by PpuMI digestion (NEB) and cassettes were serially added by Gibson assembly with PpuMI linearization of the growing array for each cycle (Primers Table 7). The final constructs were then back-cloned into the original versions of backbone and verified by analytical digestion and Sanger sequencing (Primers Table 7). The final plasmids have been validated by whole plasmid sequencing (Plasmidsaurus).

*DSB repair inhibitor treatment*

TS0111 cells were transduced with sgRNAs expressing 0-16 target sites in the human genome at MOI10. Cells were split into 96-well plates in 1:1000 dilution for clonogenicity with media containing either DMSO or 1uM of KU-0060648 (16). When cells in non-targeting controls reached full confluence, alamarBlue Cell Viability Reagent (ThermoFisher) was added and BMG POLARstar Optima microplate reader was used for fluorescence reading to assess cell viability. Excitation was set at 544nm and emission at 590nm, with a gain of 1000 and required value of 90%.
