## Supplementary material for "CRISPR-Cas9 for selective targeting of somatic mutations in pancreatic cancers": figure S: Supplementary Table S1-11.docx

**Supplementary Tables**

Table S1. Source of genomic DNA and mutation profile of the driver genes of three pancreatic cancer cases.

Table S2. Novel SVs discovered for sgRNA design.

Table S3. Novel PAMs discovered from SBSs using WGS.

Table S4. Novel PAMs discovered from SBSs using WES.

Table S5. Summary of base substitutions and somatic PAMs obtained from different ICGC projects.

Table S6. sgRNAs for growth inhibition assay.

Table S7. Number of target sites of sgRNAs for Cas9 activity and mouse-human co-culture assays in both mouse (mm10) and human (hg38) genomes.

Table S8. sgRNAs included in NT quad and Panc10.05 quad.

Table S9. TS0111-specific sgRNAs.

Table S10. Cutting efficiency and number of potential off-target sites of sgRNAs included in Panc480-MT7.

Table S11. Lowest number of mismatches of potential off-target sites in Panc480-T14 and Panc1002-T14 treated with Panc480-MT7 multiplex sgRNA expression vector.

**Table S1. Source of genomic DNA and mutation profile of the driver genes of three pancreatic cancer cases.**

| **Sample** | **Source of tumor DNA** | **Source of normal DNA** | **Tumor *KRAS*** | **Tumor *CDKN2A*** | **Tumor *SMAD4*** | **Tumor *TP53*** |
| --- | --- | --- | --- | --- | --- | --- |
| Panc480 | Primary | Lymph | G12D | Frameshift | Homozygous deletion | V274A |
| Panc504 | Primary | Duodenum | G12V | Homozygous deletion | Homozygous deletion | Frameshift |
| Panc1002 | Primary | Lymph | Q61H | Homozygous deletion | Homozygous deletion | R248Q |

**Table S2. Novel SVs discovered for sgRNA design.**

| **Cell line** | **Total no. of somatic SVs** | **No. of Sanger-validated SVs** | **No. of SVs with PAM** | **No. of good sgRNAs^#^** |
| --- | --- | --- | --- | --- |
| Panc480 | 38 | 31 | 24 | 17 |
| Panc504 | 37 | 29 | 18 | 15 |
| Panc1002 | 31 | 30 | 25 | 18 |
| **Average** | **35** | **30** | **22** | **17** |

^#^ “Good sgRNA” is defined as sgRNAs that have >50 specificity score (prediction of how much the sgRNA sequence may lead to off-target cleavage) in CRISPOR. It includes sgRNAs that are inefficient (low knockout frequencies).

**Table S3. Novel PAMs discovered from SBSs using WGS.**

| **Cell line** | **No. of SBS** | **No. of somatic PAM^&^** | **% PAM** | **No. of PAM with VAF >95%** | **No. of good sgRNAs^#^** | **No. of Sanger-validated good sgRNAs** |
| --- | --- | --- | --- | --- | --- | --- |
| Panc480 | 4576 | 385 | 8.4 | 23 | 13 | 13 |
| Panc504 | 4502 | 417 | 9.3 | 76 | 48 | 47 |
| Panc1002 | 4566 | 448 | 9.8 | 78 | 38 | 37 |
| **Average** | **4548** | **417** | **9.2** | **63** | **33** | **32** |

^&^Somatic PAM indicates a SBS of NGN/NNG sequence to NGG (both + and - strands). Only mutations with a variant allele frequency (VAF) of at least 30% in tumor (to account for subclonal mutations that potentially arose from *in vitro* culture) and a minimum of 18X read depth in both normal and tumor were included.

^#^ “Good sgRNA” is defined as sgRNAs that have >50 specificity score (prediction of how much the sgRNA sequence may lead to off-target cleavage) in CRISPOR. It includes sgRNAs that are inefficient (low knockout frequencies).

**Table S4. Novel PAMs discovered from SBSs using WES.**

| **Cell line** | **Total no. of somatic mutations** | **No. of novel PAM** | **No. of good sgRNAs^#^** | **No. of good sgRNAs with PAM of VAF >95%** |
| --- | --- | --- | --- | --- |
| Panc480 | 44 | 8 | 5 | 2 |
| Panc504 | 38 | 3 | 0 | 0 |
| Panc1002 | 30 | 4 | 2 | 0 |
| **Average** | **37** | **5** | **2** | **1** |

^#^ “Good sgRNA” is defined as sgRNAs that have >50 specificity score (prediction of how much the sgRNA sequence may lead to off-target cleavage) in CRISPOR. It includes sgRNAs that are inefficient (low knockout frequencies).

**Table S5. Summary of tumor purity, base substitutions, and somatic PAMs obtained from different ICGC projects.**

| **Project** | **N** | **% tumor purity** | | **No. of base substitutions** | | **No. of somatic PAM** | | **% PAM^*^** | |
| --- | --- | --- | --- | --- | --- | --- | --- | --- | --- |
|  |  | **Median** | **IQR**^#^ | **Median** | **IQR**^#^ | **Median** | **IQR**^#^ | **Median** | **IQR**^#^ |
| APGI-AU | 44 | 29.7 | 29.2-40.1 | 5890.5 | 4058.8-8390.3 | 478.5 | 344.8-844.0 | 8.9 | 8.1-10.5 |
| PACA-CA | 130 | 38.2 | 29.8-47.8 | 5354.5 | 4232.8-7942.0 | 430.5 | 340.5-711.5 | 8.4 | 7.7-9.8 |
| LUCA-KR | 29 | 36.3 | 30.8-47.3 | 30553.0 | 19081.5-45893.0 | 2790.0 | 2211.5-3675.0 | 8.5 | 7.8-9.2 |
| OCCAMS-GB | 388 | 32.8 | 29.5-40.0 | 20106.0 | 13542.5-31705.0 | 3235.5 | 1741.3-6167.3 | 16.1 | 12.3-20.5 |
| **All** | **591** | **34.4** | **29.5-41.0** | **15552.0** | **7091.0-26989.0** | **2131.0** | **662.0-4535.0** | **12.9** | **9.0-18.2** |

^#^IQR indicates interquartile range (25^th^-75^th^ percentile).

*% PAM = No. of somatic PAM / No. of base substitutions

**Table S6. sgRNAs for growth inhibition assay.**

| **sgRNA** | **Sequence^1^** | **Number of perfect target sites (hg38)^2^** | **Number of potential off-target sites (hg38)^2^** | **Potential target sites in exons (hg38)^3^** | **Doench ’16 predicted efficiency score^4^** | **Function** |
| --- | --- | --- | --- | --- | --- | --- |
| NT | GTATTACTGATATTGGTGGG | 0 | 0-1-12 | 0-0-0-0 | NA | Negative control |
| NT2 | GCGAGGTATTCGGCTCCGCG | 0 | 0-0-2 | 0-0-0-0 | NA |  |
| 52F(3) | TAATTACTGCACGATGCGCA | 3 | 0-0-2 | 0-0-0-0 | 59 | Multitarget sgRNAs |
| 715F(5) | ATATATATGCGATCGAGCCC | 5 | 2-1-5 | 0-0-0-0 | 54 |  |
| 551R(8) | TTGAATTGAGTTGCAACCGA | 8 | 2-1-4 | 0-0-0-0 | 61 |  |
| 230F(12) | TTGTCCCACAATGATACTTG | 12 | 8-1-8 | 0-0-0-0 | 61 |  |
| 164R(14) | GGATATTTCACTACAGACTT | 14 | 5-2-15 | 0-0-0-0 | 53 |  |
| L1.4_209F | TGCCTCACCTGGGAAGCGCA | 604 | 939-1710-2213 | NA | 55 | Positive control |
| ALU_112a | TTGCCCAGGCTGGAGTGCAG | Repeat | NA**^5^** | NA | 58 |  |

1. Sequences are followed in the genome by canonical (NGG) and/or non-canonical (NGA/NAG) PAMs. 2. CRISPOR analysis of the sgRNAs to identify the potential perfect and off-target sites (1-2-3 mismatches) in the hg38 human reference genome. 3. Number of perfect and off-target sites (0-1-2-3 mismatches) that fall within exons. 4. Cutting efficiency score based on data trained by Doench *et al.* 2016. Recommended for sgRNAs expressed with U6 promoter. The higher the efficiency score, the more likely is cleavage at this position. 5. Not applicable.

**Table S7. Number of target sites of sgRNAs for Cas9 activity and mouse-human co-culture assays in both mouse (mm10) and human (hg38) genomes.**

| **sgRNA** | **Sequence^1^** | **No. of target site in hg38^2^**  **(0-1-2-3 mismatches)** | **No. of target site in mm10^2^**  **(0-1-2-3 mismatches)** | **Function** |
| --- | --- | --- | --- | --- |
| NT2 | GCGAGGTATTCGGCTCCGCG | 0-0-0-2 | NA^4^ | Cas9 activity assay |
| HPRTc.465 | TGGATTATACTGCCTGACCA | 1-0-2-8 | NA^4^ |  |
| mOT^3^ | GGGGCTGTACTGCTTAACCA | NA^4^ | 1-0-0-10 |  |
| mchrX:52M | TATACCTAATCATTATGCCG | NA^4^ | 1-0-0-7 |  |
| NT | GTATTACTGATATTGGTGGG | 0-0-1-12 | 0-0-3-6 | Mouse-human co-cultures |
| 230F(12) | TTGTCCCACAATGATACTTG | 12-8-1-8 | 0-0-1-13 |  |

1. Sequences are followed in the genome by canonical (NGG) and/or non-canonical (NGA/NAG) PAMs. 2. CRISPOR analysis of the sgRNAs to identify the potential perfect and off-target sites (1-2-3 mismatches) in the hg38 human reference genome. 3. Off-target sgRNA. 4. NA: not applicable.

**Table S8. sgRNAs included in NT quad and Panc10.05 quad.**

| **Quad** | **Target** | **sgRNA sequence** | **PAM** | **Location type** | **Potential off-target sites (0-1-2-3 mismatches) in hg38^1^** |
| --- | --- | --- | --- | --- | --- |
| NT | NA^2^ | GGAATCATCTTCACAGTTGT | NA | NA | 0-0-4-28 |
|  | NA | AATATCCTGCCACCTCTAAC | NA | NA | 0-1-0-8 |
|  | NA | CCTCCGTCAGCAGCTAACCC | NA | NA | 0-0-0-9 |
|  | NA | ACAGATGGAGCCCAACAGAG | NA | NA | 0-0-1-35 |
| Panc10.05 | chr13:67159869 | GAGTGGCCTGTGATGACACT | GGG | *PCDH9* intronic | 0-0-1-19 |
|  | chr13:98046823 | GCAGAAAGAGATAGAATGGT | GGG | Intergenic | 0-0-3-52 |
|  | chr3:67534507 | GGGCCTTACCTGAAAGCAGC | AGG | *SUCLG2* intronic | 0-0-1-23 |
|  | chr3:76973237 | AGAATTTGAGCGACAGTATG | TGG | *ROBO2* intronic | 0-0-2-9 |

1. CRISPOR analysis of the sgRNAs to identify the potential perfect and off-target sites (1-2-3 mismatches) in the hg38 human reference genome. 2. NA: not applicable.

**Table S9. TS0111-specific sgRNAs.**

| **Pool #** | **Target** | **sgRNA sequence** | **PAM** | **Location type** | **Potential off-target sites (0-1-2-3 mismatches) in hg38^*^** |
| --- | --- | --- | --- | --- | --- |
| 1 | chr1:16152796 | AATGCTGGCTCGACAGGCTG | AGG | Intergenic | 0 - 0 - 0 - 12 |
|  | chr10:70092203 | AATTCAGTGGACGACGCCGA | GGG | PBLD intronic | 0 - 0 - 0 - 1 |
|  | chr12:10876057 | TCATTAGCATTTAAAGGCGC | CGG | Intergenic | 0 - 0 - 0 - 8 |
|  | chr12:53815822 | TCGACCCCTTCGGCCGGGCG | CGG | Intergenic | 0 - 0 - 1 - 8 |
|  | chr12:58246976 | TTCTTGAGGCCAGAACGAAG | CGG | Intergenic | 0 - 0 - 4 - 23 |
|  | chr12:92863772 | CACGTGGACAGGGCTGAAGC | CGG | LINC02397 | 0 - 0 - 4 - 23 |
|  | chr12:106150227 | AATTAGCCGGAGTGGTGGTG | GGG | Intergenic | 0 - 0 - 33 - 48 |
|  | chr12:128055569 | ACATGGTGCCCCGTCGGCTA | CGG | Intergenic | 0 - 0 - 0 - 2 |
|  | chr12:130765849 | TGGGCCCAGGCTCGGGGGCT | GGG | Intergenic | 0 - 0 - 7 - 50 |
| 2 | chr14:30809300 | AATCATGATGTCTGTCTTCA | TGG | Intergenic | 0 - 0 - 3 - 63 |
|  | chr14:39901580 | CAGCGGCCCGGAAGCCTCAA | GGG | FBXO33 UTR | 0 - 0 - 0 - 5 |
|  | chr14:42234257 | TTTCAAGACGTTAAAGAAAC | AGG | LRFN5 intronic | 0 - 0 - 5 - 33 |
|  | chr14:69405738 | GCCAAGAAGCAGGGGGCCTG | CGG | ACTN1 intronic | 0 - 0 - 7 - 39 |
|  | chr14:75084005 | TTTGGAAGGTGCAGGCCGTA | CGG | Intergenic | 0 - 0 - 1 - 3 |
|  | chr14:75780274 | CCACAAAGTACACAAAGAAC | AGG | Intergenic | 0 - 0 - 5 - 23 |
|  | chr14:91885563 | TTAATTGCTTCTCCGCCCGC | CGG | Intergenic | 0 - 0 - 0 - 1 |
|  | chr16:14136787 | GGCTTTGTTTATGGGACAGA | TGG | Intergenic | 0 - 0 - 3 - 21 |
|  | chr16:55468556 | AACCCCGGCCACCCAGGCTG | GGG | MMP2-AS1 intronic | 0 - 0 - 13 - 106 |
| 3 | chr16:58087434 | ACGTGGTCAGCCACCACAGA | CGG | Intergenic | 0 - 0 - 1 - 14 |
|  | chr16:75121047 | AGAGTTCTTGTAGCTTGAAC | CGG | ZNRF1 intronic | 0 - 0 - 2 - 14 |
|  | chr16:78133950 | GACCCGAGGCCCACCAAGGG | GGG | WWOX UTR | 0 - 0 - 1 - 17 |
|  | chr16:79964797 | AGGCCAAGTGGATGGATGAT | GGG | Intergenic | 0 - 0 - 4 - 25 |
|  | chr16:89336911 | GTGAGCCCCCTCTAAGGTCC | CGG | ANKRD11 intronic | 0 - 0 - 2 - 5 |
|  | chr17:29776374 | TCCTTTCCATTCGCTTCTGG | AGG | RAB11FIP4 intronic | 0 - 0 - 0 - 9 |
|  | chr17:38082034 | GCATCTCTTCAATCAGAATG | CGG | ORMDL3 exonic | 0 - 0 - 1 - 22 |
|  | chr17:44928421 | AACCGCTCTCTGGAAGCGGG | GGG | Intergenic | 0 - 0 - 1 - 3 |
|  | chr17:71383391 | ATTTAAAAAAGAGAGGCCAG | GGG | SDK2 intronic | 0 - 0 - 15 - 75 |

^*^CRISPOR analysis of the sgRNAs to identify the potential perfect and off-target sites (1-2-3 mismatches) in the hg38 human reference genome.

**Table S10. Cutting efficiency and number of potential off-target sites of sgRNAs included in Panc480-MT7.**

| **Target** | **sgRNA sequence** | **PAM** | **Mutation type (copy number)** | **Mutation frequency (%)^&^** | **No. of potential off-targets (0-1-2-3-4mm)^*^** | **No. of potential off-targets including NAG PAM^$^** |
| --- | --- | --- | --- | --- | --- | --- |
| chr8:201457 | GGAATCATCTTCACAGTTGT | TGG | D-LOH^#^ (1) | 22.6 | 0-0-3-20-159 | 1-0-3-44-376 |
| chr17:5377742 | AATATCCTGCCACCTCTAAC | AGG | D-LOH (1) | 36.4 | 0-0-0-6-96 | 1-1-0-16-234 |
| chr3:537601 | TCAGTCCAGTCAAAGGTGGA | AGG | D-LOH (1) | 87.3 | 0-0-1-7-119 | 0-0-3-26-281 |
| chr3:59525282 | CTAATGTATGACTGAAAGCT | GGG | D-LOH (1) | 71.1 | 0-0-1-7-137 | 0-0-1-22-359 |
| chrX:3982448 | GAGGTGTCTAAACCATGACA | AGG | D-LOH (1) | 67.8 | 0-0-0-4-103 | 1-0-1-15-230 |
| chr8:29032916 | GTGCACATCTTATCTCCCTT | AGG | D-LOH (1) | 57.6 | 0-0-0-8-117 | 0-1-1-14-271 |
| chr18:1819017 | TTAGGGGGCCAAGAGCGTAT | GGG | D-LOH (1) | 68.7 | 0-0-0-2-32 | 0-0-0-5-67 |

^#^D-LOH: deletion-based loss of heterozygosity

^&^Individual sgRNAs were transduced into Panc480 cells separately and puromycin-selected for 7 days. Cells were harvested for NGS and mutation frequency was quantified using CRISPResso2.

^*^sgRNA sequences were put through Cas-OFFinder to identify potential off-target sites with 1-4 mismatches in hg19. Only sites with the canonical NGG PAM were included.

^$^sgRNA sequences were put through Cas-OFFinder to identify potential off-target sites including ones with non-canonical NAG PAM and ones with 1-4 mismatches.

**Table S11. Lowest number of mismatches of potential off-target sites in Panc480-T14 and Panc1002-T14 treated with Panc480-MT7 multiplex sgRNA expression vector.**

| **Target** | **sgRNA sequence** | **PAM** | **Lowest number of mismatches in Panc480-T14 replicate 1^*^** | **Lowest number of mismatches in Panc480-T14 replicate 2^*^** | **Lowest number of mismatches in Panc1002-T14^*^** |
| --- | --- | --- | --- | --- | --- |
| chr8:201457 | GGAATCATCTTCACAGTTGT | TGG | 7 | 6 | 7 |
| chr17:5377742 | AATATCCTGCCACCTCTAAC | AGG | 6 | 7 | 7 |
| chr3:537601 | TCAGTCCAGTCAAAGGTGGA | AGG | 7 | 7 | 7 |
| chr3:59525282 | CTAATGTATGACTGAAAGCT | GGG | 5^#^ | 6 | 6 |
| chrX:3982448 | GAGGTGTCTAAACCATGACA | AGG | 6 | 7 | 7 |
| chr8:29032916 | GTGCACATCTTATCTCCCTT | AGG | 4^#^ | 7 | 6 |
| chr18:1819017 | TTAGGGGGCCAAGAGCGTAT | GGG | 6 | 8 | 7 |

*WGS analyses were performed for T14s. For each indel detected by Mutect2, the original sequence on the reference genome was compared to the sgRNA sequence to determine the homology between both using the Smith-Waterman algorithm (see Methods). The lowest number of sequence mismatch was shown. On-target mutation was excluded in this analysis as results had been shown in figure S5B using a deep sequencing assay.

^#^One mutation only.
